# High-fidelity but hypometric spatial localization of afterimages across saccades

**DOI:** 10.1101/2025.05.04.652081

**Authors:** Richard Schweitzer, Thomas Seel, Jörg Raisch, Martin Rolfs

**Affiliations:** Centro Interdipartimentale di Mente e Cervello, Università degli studi di Trento, Italy; Department of Psychology, Humboldt-Universität zu Berlin, Germany; Science of Intelligence, Research Cluster of Excellence, Germany; Control Systems Group, Technische Universität Berlin, Germany; Institute of Mechatronic Systems, Leibniz Universität Hannover, Germany; Bernstein Center for Computational Neuroscience Berlin, Germany

**Keywords:** complete darkness, afterimages, sensorimotor contingencies, saccades, spatial localization, efference copy, visual stability

## Abstract

Humans typically perceive their visual world as stable and continuous, despite frequent shifts of the retinotopic reference frame caused by saccades. This visual stability is paralleled by afterimage movement across saccades: Although retinotopically stable, afterimages appear to move in egocentric space wherever the eye moves. To investigate the mechanisms underlying this phenomenon, we tasked human observers to localize afterimages relative to briefly flashed probes in complete darkness. This psychophysical tracking of afterimages was accompanied by eye tracking, allowing us to fit a dedicated computational model to accurately predict afterimage movement based on the size of eye movements. The gain of afterimage movement was significantly hypometric, remained unaffected by post-saccadic visual feedback and saccadic adaptation, and was inversely related to saccade gain. These findings suggest that afterimage movement is driven by efference-based, feedforward prediction of visual consequences of saccades and demonstrate the potential of the afterimage-tracking technique for studying visual stability.

**Significance:** In low-light environments, brief high-intensity visual stimulation can induce long-lasting retinal afterimages. When observers then make eye movements to explore their visual environment, these afterimages – albeit fixed in the retinotopic frame of reference – appear to move in egocentric space wherever the eye moves. Even though this phenomenon has been known for centuries, the underlying computations remained unexplained. Tracking eye and afterimage positions simultaneously, we found that perceived afterimage position was accurately predicted by eye position across a variety of visuomotor conditions, whereby the eye movement’s size was however systematically underestimated by the visual system. Considering a parsimonious model of visual localization, afterimage movement can be understood as a consequence of feedforward predictions of the visual consequences of impending eye movements.

## Introduction

When active observers make rapid eye movements – saccades – to visually explore their environment, each movement is associated with an inevitable shift of the retinal image in the opposite direction. Despite the constant repositioning of the retinotopic frame of reference, the egocentric view of the world remains stable and continuous– a phenomenon known as visual stability. The question of how this amazing feat is realized by visual system has been addressed by countless investigations (for reviews, see, e.g., Bridgeman, Van der Heijden, & Velichkovsky, 1994; Cavanagh, Hunt, Afraz, & Rolfs, 2010; MacKay, 1972, 1973; Melcher, 2011; Rolfs, 2015; Wurtz, 2008), studies have converged to explanations that invoke senso-rimotor predictions based on corollary-discharge (Sperry, 1950) or efference-copy (Von Holst & Mittelstaedt, 1950) signals that are grounded in known neuronal structures and pathways (Sommer & Wurtz, 2008; Wurtz, 2018). Whereas their relevance is undeniable, still little is known about the accuracy of such efference-based sensorimotor predictive mechanisms.

A promising venue to understand how well the visual system can account for the consequences of saccades, is the curious case of phenomenological movement of retinal afterimages in egocentric visual space. While objects whose projections shift across the retina due to saccades appear stable, the opposite is the case for afterimages: Once induced, they remain in a fixed retinotopic location but move in space contingently upon the eye movements one makes. When observing afterimages in complete darkness, that is, in the absence of any visual input, their movement can only be created by relying on non-visual sources of information. Proprioceptive information from the eye muscles’ stretch receptors, on the one hand, has been considered unlikely to account for this movement: Whereas briefly pressing one’s eye ball while occluding the other eye results in movement of the entire visual world – a disruption that proprioceptive information is apparently unable to compensate for (Bridgeman et al., 1994) – the spatial position of an afterimage remains unaltered (e.g., Grüsser, 1994; Mittelstaedt, 1990). Active eye movements, on the other hand, produce reliable movement of afterimages, making motor efference the most likely source to inform subjective afterimage position. Even though this has been known for more than two centuries and tracking afterimages has even been used as an early eye-tracking technique (Wade, 2015), the exact relationship between saccade and resulting afterimage movement is poorly understood. Using saccades to auditory cues rapidly alternating between two locations, Grüsser, Krizic, and Weiss (1987) estimated that the spatial extent of afterimage movement amounts to merely 50-85% of the saccade amplitude – a result well compatible the efference-copy gain of 0.625 estimated by Bridgeman and Stark (1991) – which was even further reduced as saccade rate increased. Given such significant underestimation of the size of actual eye movements, it is unclear to what extent sensory predictions informed by this information could contribute to obtaining visual stability.

Here, we assessed the extent of afterimage movement across saccades and concurrently registered gaze position. We discovered a remarkable congruence of eye and afterimage movements with gains above 0.9, yet significantly below 1, which was independent of saccade direction and amplitude, post-saccadic visual references, and even a systematic reduction of saccade amplitudes using saccadic adaptation. These remarkable results suggest that to realize accurate movement of retinotopically static stimuli in egocentric visual space, the oculomotor system is capable of generating accurate but hypometric approximations of the sensory consequences of its own saccades.

## Results

Phenomenological movement of afterimages can be readily observed in low-light environments after fixating a bright source of light for a few seconds. After the light source is extinguished, a localized afterimage remains that shifts in space according to where gaze is shifted. While the phenomenon is easy to experience, it is harder to study in an experimentally controlled setting. To this end, we developed a custom array of LEDs to be used for visual stimulation in an otherwise completely dark environment (see Apparatus). To allow for sufficient time for dark adaptation, as well as to evaluate the functionality of the apparatus with a more conventional paradigm, all experimental sessions started with a pre-test featuring a trans-saccadic target-displacement discrimination task entirely unrelated to subsequent afterimage localization (for details, see Pre-experiment). In this pre-test, we estimated the post-saccadic perceptual null location (PNL) of saccade targets in a range of timing conditions (Figure A1a,b) and found that saccade targets had to be shifted by up to 2 degrees of visual angle (dva) in the direction of the saccade to be perceived as stationary (Figure A1c–e). Together with the result that this tendency was strongest when saccade targets were extinguished during saccades, this finding provides preliminary evidence in favor of the hypothesis that visual persistence of the target stimulus, that is, the extension of activation beyond the time of stimulus offset due to the properties of the temporal response function (Bowen, Pola, & Matin, 1974), can well lead to the impression of it moving along with the executed eye movement. Notably, this hypothesis has already been put forward to explain peri-saccadic flash mislocalization (Pola, 2004; Teichert, Klingenhoefer, Wachtler, & Bremmer, 2010) and is compatible with the saccade-contingent movement of afterimages: Both visual persistence and afterimages are fixed in retinotopic space (Duysens, Orban, Cremieux, & Maes, 1985) and should – following the reasonable assumption that efference-copy signals are employed to perform spatial localization across saccades – be expected to move in the direction of saccades in egocentric space. Given the considerably longer durations of afterimages (Robertson & Fry, 1937), their amplitude of perceived movement should be proportionately larger, making them an ideal stimulus to study motorto-visual transformations based on efference-copy signals.

Upon sustained dark adaptation (i.e., about 10-20 min after starting the experimental session), we switched to a novel psychophysical localization paradigm in which observers localized afterimages relative to visual probes briefly flashed in total darkness across even sequences of saccades. As illustrated in Figure 1a, afterimages were induced centrally by fixation of a repetitively flashed, very bright pulse, which was directly followed by the brief presentation of the saccade target that instructed the primary saccade. After the execution of primary saccades, observers indicated when they had a clear view of the afterimage and thereby triggered a sequence of localization probes. Note that this step was necessary as afterimages need time not only to be generated, but also to emerge again after saccades (for a review on saccadic suppression of afterimages and entoptic images, see Matin, 1974). Afterimages were localized with respect to the location of each of up to seven localization probes (for details, see Main experiment). This approach allowed us to take into account changes of afterimage position across multiple eye movements.

**Figure 1.**
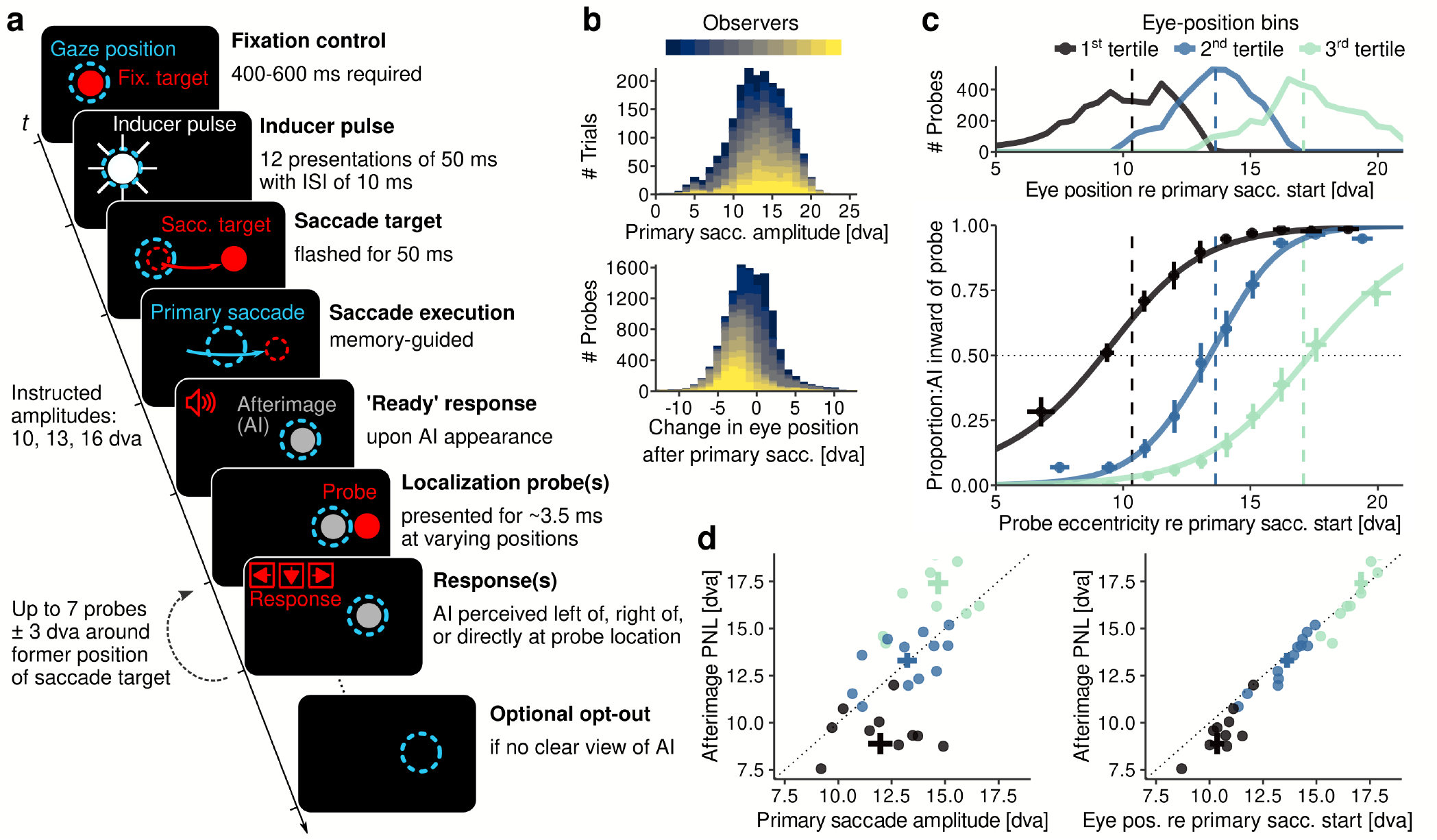
Tracking afterimages across saccades (Session 1). **a** Illustration of experimental paradigm used to localize an afterimage (AI) in complete darkness (stimuli not drawn to scale). For details, see main text and Main experiment. **b** Histograms of primary saccade amplitude (top) and eye position at the time of localization-probe presentation relative to the primary saccade’s landing position (bottom). Colors indicate individual observers. **c** Results of the binning of eye position at the time of probe presentation relative to each trial’s corresponding primary saccade’s start position (top) and resulting average psychometric functions (bottom). Colors indicate the three eye-position bins and vertical dashed lines their average value. **d** Individual observers’ afterimage PNLs as a function of primary saccade amplitude (left) and eye position (right). Crosses represent mean ±1 SEM values in each eye-position bin.

### Afterimage movement across saccade amplitudes

As a first step, we quantified the amplitude of afterimage movement relative to a wide range of saccade amplitudes (Session 1: Saccade amplitude). Primary saccade amplitudes amounted to 13.34 dva (SD = 1.65) and exhibited considerable variance within each participant (mean SD: 3.06 dva; Figure 1b, top). However, even after primary saccades, the eyes continued to move considerably in complete darkness (mean SD: 2.39 dva; Figure 1b, bottom), underscoring the necessity of considering postsaccadic eye movements to correctly compute afterimage position. We thus mapped localization-probe positions and eye positions at the time of probe presentation to a common scale with reference to each primary saccade’s onset. To aggregate localization judgments across probe and eye positions and estimate afterimage PNLs, we discretized the scale by binning, individually for each participant, and then fitted psychometric functions (see Analyses). Aggregated results revealed a clearly discernible pattern: Afterimage PNLs varied significantly with the eyes’ displacement relative to the start position of the primary saccade, that is, the position where the afterimage was induced (*F* (2, 22) = 205.64, *η*^2^ = 0.80, *p* < .001, *p_GG_ <* .001; Figure 1c). To investigate whether the variance of afterimage PNLs scaled in a linear fashion with eye-movement amplitude, we ran separate linear regressions on primary saccade amplitudes and eye position to predict the position of afterimages. While primary saccade amplitude significantly predicted PNLs (*β* = 1.19, *t* = 4.64, *p* < .001; Figure 1d, left), the differences in saccade amplitude did not align too well with afterimage po-sitions 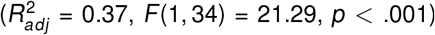. In contrast, performing the same analysis for eye position at the time of probe presentation yielded a near-perfect prediction of afterimage PNLs (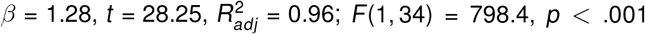; Figure 1d, right), which remained highly significant even when the variance introduced by binning was removed by subtracting the mean eye-movement amplitude in each bin 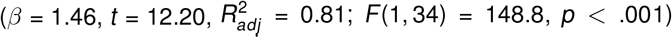. These comparisons make clear that accounting for eye movements performed after primary saccades is critical: Even in total darkness, where no post-saccadic visual references are available, secondary saccades may well be executed according to an efference-based error signal (Ohl, Brandt, & Kliegl, 2013). Indeed, as evidenced by Figure 1d and estimated regression slopes above 1, primary saccades seemed to overshoot when instructed amplitudes were short and to undershoot when they were large (left panel), yet eye positions were already corrected when probes were presented (right panel). It should be noted, however, that even with this correction, afterimage positions were not perfectly aligned with eye positions, providing a first intuition that the impressive congruence between eye and afterimage positions may be subject to systematic errors.

### Modeling head-centered afterimage position

To account for the large variance in eye-movement behavior across observers and trials, especially with respect to eye movements performed after primary saccades, we developed a computational model (henceforth, the Gain Model) to track afterimage position relative to eye position over time. To elaborate, previous analyses assumed the independence of individual eye positions when predicting the subjective location of the afterimage – an assumption that neglects that afterimage position is most likely a function of an entire sequence of eye movements that have occurred up to the presentation time of each individual localization probe. These eye movements may encompass saccades, microsaccades, ocular drift, or even pursuit of the afterimage. Note that attempts to fixate slightly peripheral afterimages inevitably fail due to the their gazecontingent movement, and there is good evidence that a high-resolution eye-position signal is available even for drift motion (Zhao, Ahissar, Victor, & Rucci, 2023). As afterimage movement was clearly observed as a consequence of all eye movements, the following analyses parsimoniously treat all types of eye movements alike by considering only their resulting gaze position at the time of probe presentation. This approach allows for a characterization of afterimage movement that not only goes beyond the effect of the primary saccade, but also accounts for (potentially large) oculomotor variability across trials and observers.

The Gain Model, outlined in Figure 2a-b (for further details, see Estimation of afterimage movement gain), states that the (unknown) spatial location of the afterimage at a given time point (denoted *x_i_*) changes proportionally to the size of each change in eye position (that is, *g_i_* − *g*_*i*−1_) that occurs between the presentations of probes. Note that, in order to implement this correctly, a starting value *x*_0_ was needed: After correcting for the estimated retinal eccentricity of the afterimage (Figure B2), the afterimage could be assumed to be approximately foveal. Critical to the model, a (unknown) sensorimotor gain *G* translates the change in eye position to a change in afterimage position. If one assumed that afterimage position is perfectly contingent upon eye position (i.e., G=1), then eye and afterimage positions would be equal. To fit the Gain Model to the data, a response mechanism was implemented that was based on the relative horizontal distance between (modeled) afterimage position and (recorded) probe position (Figure 2b). As observers had the option to report afterimages ‘directly at the probe location’, we included a range *r* within which the afterimages would be judged as in the same position as the probe, and to model potential response biases, we also included a center variable *m*, around which range *r* was located. To estimate parameters *G, m*, and *r*, we performed a grid search and selected those sets of parameters that best approximated each observers’ responses. To evaluate how well model predictions generalized over primary-saccade directions, grid-search simulations were run separately on trials with only rightward or leftward primary saccades, as well as on all trials including both directions. Figure 2c shows that, when fitted on the entire set of a single observer’s data, the Gain Model was able to correctly classify 84.8% (SD = 4.07, range [77.31, 91.16]) of responses. Since fitting separately for primary-saccade directions neither improved classification accuracy (*F* (2, 22) = 1.84, *η*^2^ = 0.01, *p* = .182, *p_GG_* = .201) nor affected parameter estimates (*G*: *F* (2, 22) = 3.11, *η*^2^ = 0.06, *p* = .065, *p_GG_* = .070; *m*: *F* (2, 22) = 1.14, *η*^2^ = 0.03, *p* = .335, *p_GG_* = .314; *r* : *F* (2, 22) = 0.96, *η*^2^ = 0.01, *p* = .399, *p_GG_* = .389), as well as for the sake of parsimony, we henceforth only considered model predictions fitted on all trials, irrespective of primary-saccade direction.

**Figure 2.**
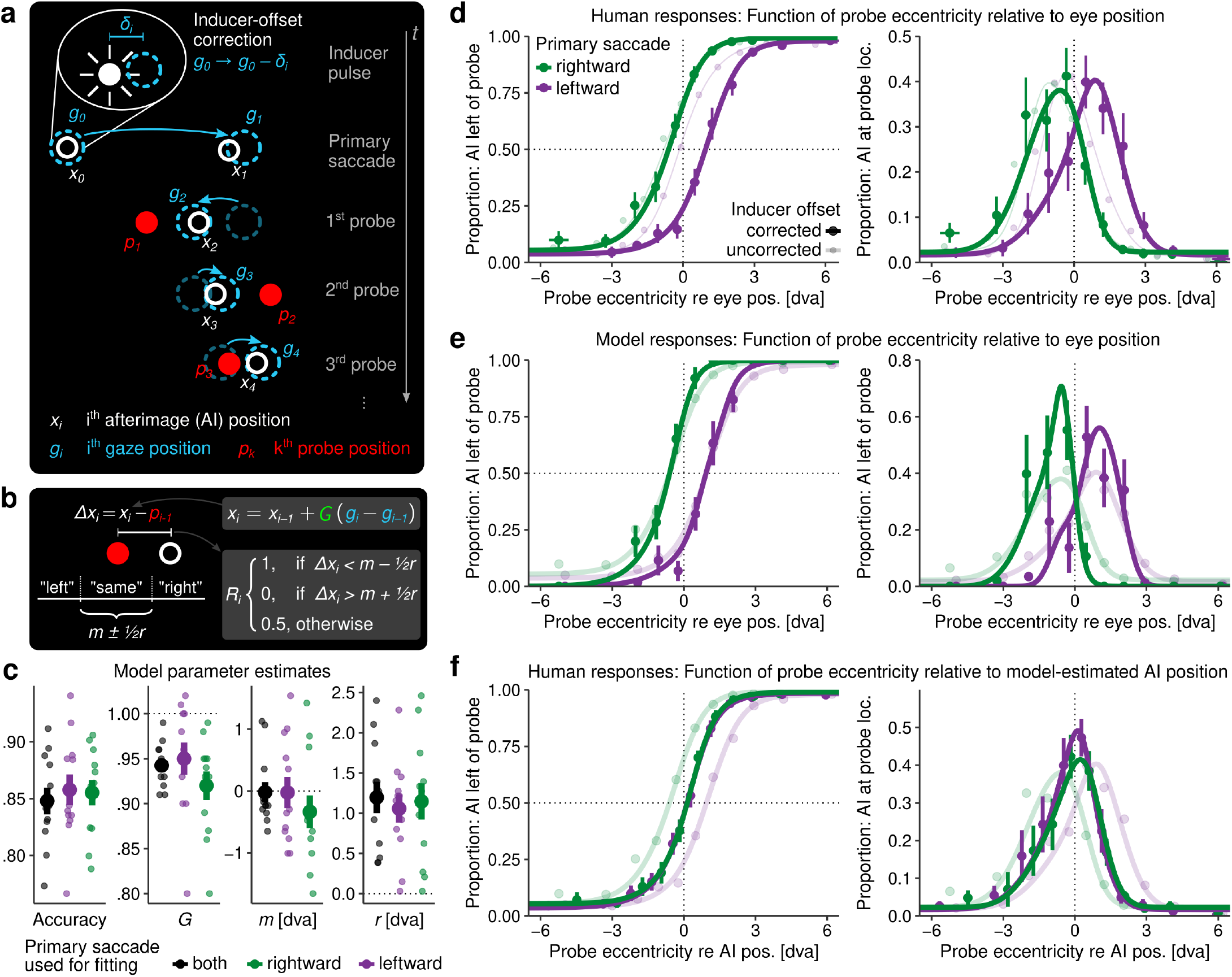
Modeling eye movement-contingent afterimage position with the Gain Model. **a** Illustration of the assumptions of the Gain Model. While gaze positions (g, in blue) and probe positions (p, in red) are known in each afterimage-localization trial, the goal is to estimate the unknown afterimage position (x, in white). All elements are drawn in a subjective egocentric (as opposed to retinotopic) frame of reference. **b** Description of model parameters and response mechanism assumed to fit the model to human localization judgments (for details, see main text and Estimation of afterimage movement gain). **c** Classification accuracy (leftmost panel) and best-fitting model parameters for each observer, for fits performed across primary-saccade directions (black dots) and separately for leftward and rightward primary saccades (purple and green dots, respectively). **d** Group-level psychometric functions, logistic (left) and Gaussian (right), approximating human localization judgments based on probe position relative to eye position. Thin transparent lines indicate results without correction for afterimage eccentricity (Figure B2). **e** Same as above, but showing instead responses of the Gain Model. Transparent lines reproduce human responses presented in panel d as reference. **f** Human localization judgments as a function of probe position relative to modeled afterimage position (x) with the same reference as in panel e.

We first analyzed to what extent afterimage localization could be understood purely in terms of retinal eccentricity of localization probes, that is, probe position relative to eye position at the time of probe presentation (Figure 2d): Both directional (left) and ‘directly-at’ (right) localization judgments revealed PNLs offset to the left for rightward saccades (Logistic: *M* = −0.63 dva, *SD* = 0.61, *t* (11) = −3.61, *p* = .004; Gaussian: *M* = −0.66 dva, *SD* = 0.74, *t* (11) = −3.12, *p* = .010) and to the right for leftward saccades (Logistic: *M* = 0.90 dva, *SD* = 0.68, *t* (11) = 4.59, *p* < .001; Gaussian: *M* = 0.74 dva, *SD* = 0.82, *t* (11) = 3.14, *p* = .009). The direction-dependent offset of psychometric functions represents an important finding, as it shows that the localization of afterimages is not performed in a purely retinotopic frame of reference. If it were, localization responses would merely depend on the retinotopic offset between the (corrected-to-foveal) afterimage and localization probe. If such a mechanism were in place, then midpoints of psychometric functions shown in Figure 2d would be at zero – which is not the case. Instead, the significant and remarkably symmetric offsets of PNLs for rightward and leftward primary saccades indicate that afterimage position cannot be equated with eye position at the time of probe presentation. In fact, the amplitude of afterimage movement systematically falls short of the actual eye movement. These patterns were reproduced by model responses with high fidelity (Figure 2e), even though logistic slopes were steeper and Gaussian amplitudes larger than in the original data because the response mechanism of the Gain Model was – unlike human response behavior – deterministic. Critically, fitted model parameters corroborated psychophysical results: *G* amounted to 0.94 (SD = 0.02, range [0.91, 0.99]), a value significantly below 1 (*t* (11) = −8.11, *p* < .001) and therefore hypometric estimate. Response parameters *m* and *r* amounted to −0.02 dva (*SD* = 0.57) and 1.20 dva (*SD* = 0.68), respectively Figure 2c). Finally, if the proposed model represented a suitable estimate for the latent perceived position of afterimages, then veridical psychometric functions would be expected when modeled afterimage positions are used to predict human responses. This was precisely what we observed: Using probe eccentricity relative to (modeled) afterimage position as a predictor (Figure 2f), logistic PNLs (left panel) of the two primary-saccade directions became indistinguishable not only from each other (*F* (1, 11) = 0.10, *η*^2^ *<* 0.01, *p* = .756) but also from zero (*F* (1, 11) = 0.41, *η*^2^ = 0.03, *p* = .532). The same was the case for PNLs estimated by the Gaussian function (intercept: *F* (1, 11) = 0.01, *η*^2^ *<* 0.01, *p* = .918; main effect primary direction: *F* (1, 11) = 2.70, *η*^2^ = 0.02, *p* = .128). In addition, as compared to using probe eccentricity as predictor, localization accuracy improved when afterimage position was used as predictor: Steeper slopes of logistic functions (*F* (1, 11) = 5.99, *η*^2^ = 0.02, *p* = .032) and reduced standard deviations of Gaussian functions (*F* (1, 11) = 5.44, *η*^2^ = 0.01, *p* = .040) were observed, suggesting that human localization responses were indeed better explained by afterimage position than by eye position. These results provide clear evidence against the alternative hypothesis that afterimages could be localized relative to the retinotopic coordinates of probes without taking into account extra-retinal information.

One central assumption of the Gain Model is that the change in afterimage position linearly scales with the change in eye position. It follows that the absolute size of the spatial discrepancy between eye and afterimage position should increase as the size of the eye movement increases. To critically validate this assumption, we first quantified the eye movement’s size (that is, eye position relative to the starting point of the primary saccade) when probes were presented. Second, as above, we computed the probe’s retinal eccentricity (that is, probe position relative to gaze position) and fitted psychometric functions separately for three eye movement-size bins (1st bin: *M* = 10.1 dva, *SD* = 1.07; 2nd bin: *M* = 13.18 dva, *SD* = 0.73; 3rd bin: *M* = 15.94, *SD* = 0.67). Figure 3a shows that afterimage PNLs were again shifted left for rightward saccades and right for leftward saccades but, most critically, the absolute size of this shift depended on the size of the eye movement (1st bin: *M* = 0.61 dva, *SD* = 0.24; 2nd bin: *M* = 0.80 dva, *SD* = 0.34; 3rd bin: *M* = 1.08, *SD* = 0.38; *F* (2, 22) = 30.62, *η*^2^ = 0.11, *p* < .001, *p_GG_ <* .001). PNLs were similar across primary saccade directions (*F* (1, 11) = 0.67, *η*^2^ = 0.04, *p* = .429) and their dependency on eye-movement size remained also largely unaltered by primary saccade direction (*F* (2, 22) = 3.11, *η*^2^ = 0.01, *p* = .064, *p_GG_* = .069). Furthermore, we found a significant linear relationship between AI PNLs (collapsed across primary saccade directions) and the corresponding eye movement’s size (*β* = 0.07, *t* = 3.38, *p* = .001, 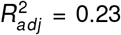; Figure 3b) with an intercept not significantly different from zero (*β* = − 0.13, *t* = − 0.45, *p* = .654). Note that the estimated slope of 0.07 was strikingly well in line with the results provided by Gain Model: The estimated average gain of 0.94 (cf. Figure 2c) would have predicted a hypometric offset between afterimage and eye movement amounting to 6% of the eye movement’s size. These results suggest that observed PNLs were not merely a product of a (direction-specific) retinotopic bias in afterimage localization, but were directly linked to the executed eye movement. They furthermore support the validity of the assumption of linear gains and thus underline the suitability of the Gain Model as a tool to describe afterimage movement across saccades in egocentric space. The model not only approximates and explains localization responses well enough, but also estimates plausible, hypometric gains by which efference-copy signals are forwarded to anticipate visual changes induced by saccades.

**Figure 3.**
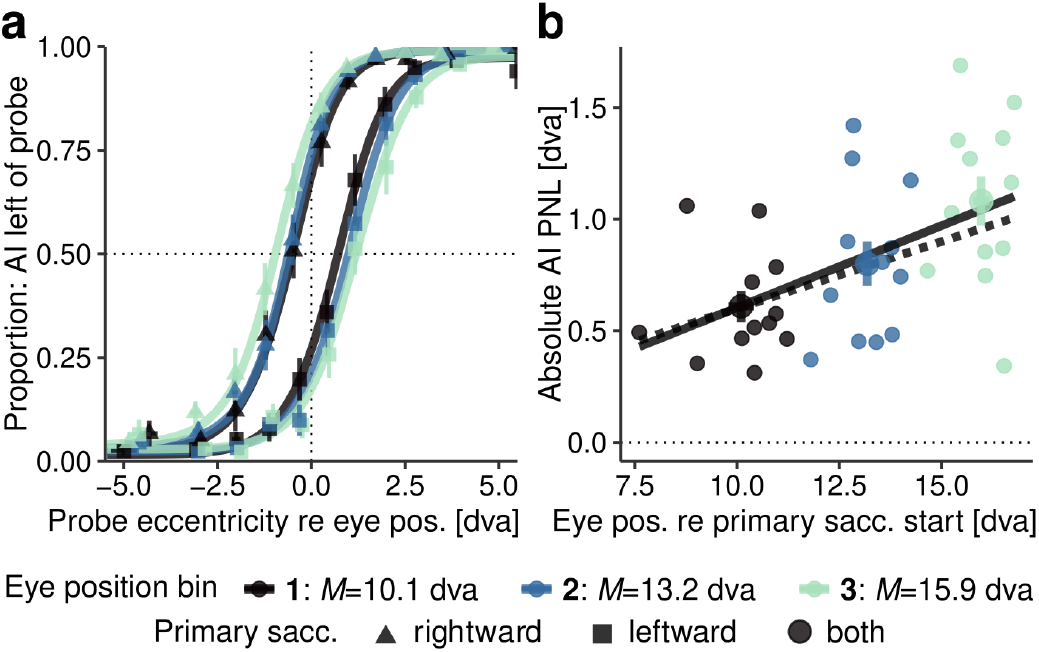
Validation of the Gain Model. **a** Group-level logistic psychometric functions approximating localization judgments in three eye-position bins. Triangles and squares show aggregated response proportions for rightward and leftward primary saccades, respectively. **b** Absolute afterimage PNLs for each observer and eye-position bin as a function of the size of the eye movement (computed relative to the starting position of the primary saccade). Values were collapsed across primary saccade directions. The dotted black line indicates the prediction based on an afterimagemovement gain of 0.94, while the solid black line shows the linear fit of the data.

### Independence of post-saccadic visual references

While the prediction of saccade-induced visual consequences is generally assumed to rely on efference copy signals, visual references in general (Deubel, 2004), but especially the saccade target itself (McConkie & Currie, 1996), serve as feedback to determine the accuracy of the performed saccade and to drive saccadic adaptation (Cassanello, Ostendorf, & Rolfs, 2019; McLaughlin, 1967). Making use of the Gain Model described above, we thus proceeded to investigate the potential role of postsaccadic visual references in the localization of afterim-ages in a second experimental session. As illustrated in Figure 4a, three types of visual feedback were administered in afterimage-localization trials (for details, see Session 2: Visual references): No feedback (as in Session 1) where the saccade landed in total darkness, normal (that is, veridical) feedback where the original pre-saccadic target remained on the screen for 150 ms, and false feedback where the pre-saccadic target was shown inward-displaced by 2 dva for the same duration. If visual feedback were used to determine afterimage position, then significant shifts of PNLs – probably in the direction of post-saccadic targets – relative to the no-feedback condition would be assumed, resulting in a modulation of the estimated gain *G*.

**Fig. 4.**
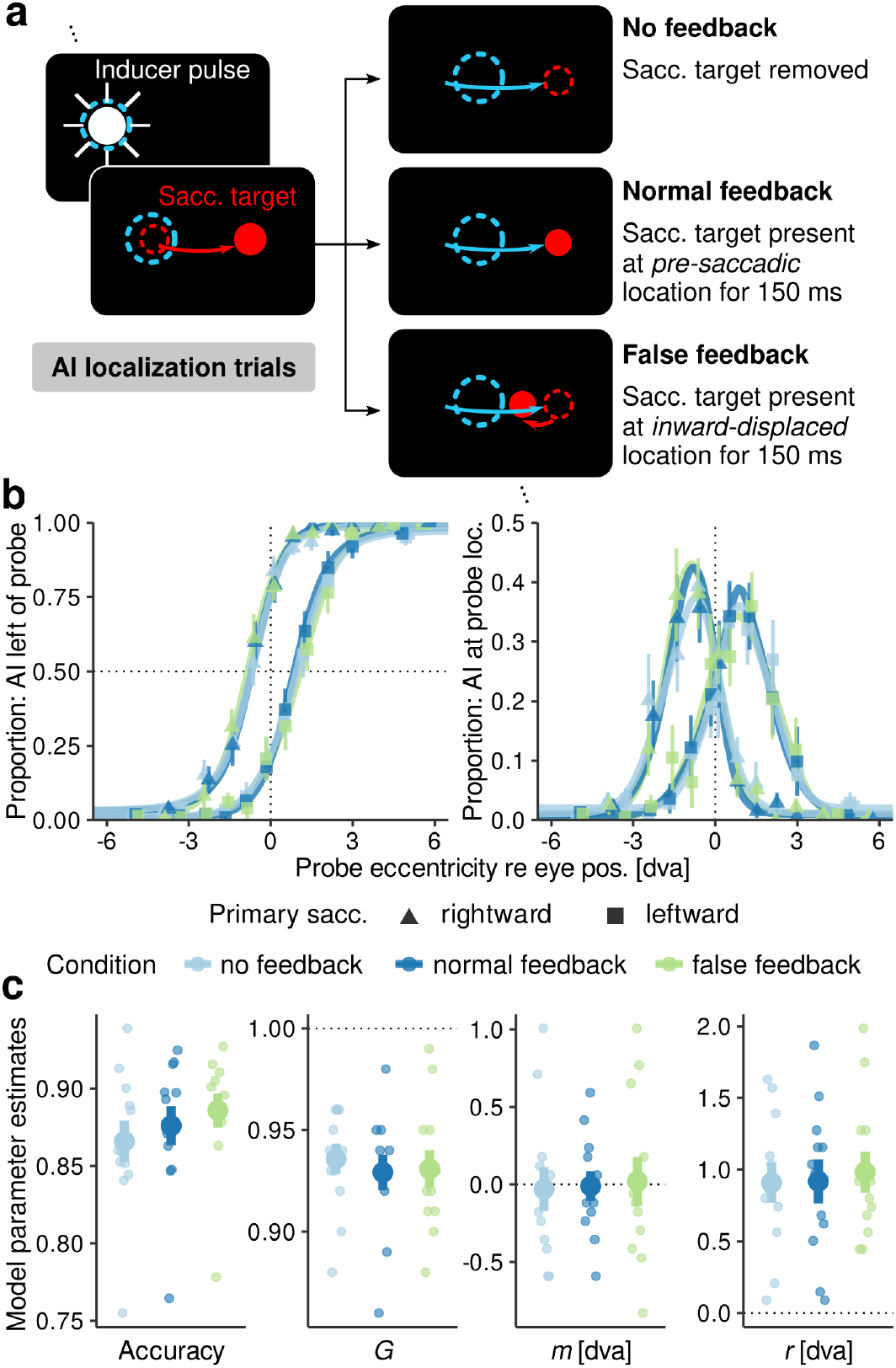
Afterimage (AI) position and post-saccadic visual feedback (Session 2). a Illustration of the three visualreference manipulations, which were all performed strictly during primary saccades (Session 2: Visual references).
b Group-level psychometric functions, logistic (left) and Gaussian (right), approximating localization judgments in
the three visual-reference conditions based on probe position relative to eye position. Triangles and squares show aggregates for rightward and leftward primary saccades, respectively. c Classification accuracy (leftmost panel) and best-fitting model parameters for each observer and condition together with group means (± SEM). All fits were performed across primary-saccade directions.

Performing the same analyses as above, this time separately for all visual-feedback conditions, we first replicated the direction-dependent PNL offsets from zero in the no-feedback condition (rightward: *M* = −0.73, *SD* = 0.60, *F* (1, 11) = 27.87, *η*^2^ = 0.68, *p* < .001; leftward: *M* = 0.92, *SD* = 0.69, *F* (1, 11) = 25.67, *η*^2^ = 0.67, *p* < .001;Figure 4b, left), suggesting a stability of these patterns across sessions. Second, we found no evidence for different PNLs in the three visual-feedback conditions (superimposed psychometric functions in Figure 4b), neither with logistic functions (rightward: *F* (2, 22) = 2.55, *η*^2^ = 0.03, *p* = .100, *p_GG_* = .119; leftward: *F* (2, 22) = 1.08, *η*^2^ = 0.01, *p* = .356, *p_GG_* = .352) nor with Gaussian functions (rightward: *F* (2, 22) = 0.20, *η*^2^ *<* 0.01, *p* = .820, *p_GG_* = .803; leftward: *F* (2, 22) = 0.09, *η*^2^ *<* 0.01, *p* = .905, *p_GG_* = .893). Finally, estimated model parameters (Figure 4c) also did not indicate any differences between conditions (*G*: *F* (2, 22) = 0.54, *η*^2^ = 0.01, *p* = .591, *p_GG_* = .583; *m*: *F* (2, 22) = 0.17, *η*^2^ *<* 0.01, *p* = .841, *p_GG_* = .825;*r* : *F* (2, 22) = 0.20, *η*^2^ *<* 0.01, *p* = .823, *p_GG_* = .811). A small but significant effect was found on the model’s classification accuracy (*F* (2, 22) = 4.37, *η*^2^ = 0.04, *p* = .025, *p_GG_* = .031; Figure 4c, leftmost panel). This unexpected effect may be interpreted in terms of improved stabilization of gaze position with post-saccadic feedback, which positively affect the model’s fit to the recorded data. In sum, the localization of afterimage appeared to be largely independent of whether brief post-saccadic visual references were available or not, providing evidence that movement of afterimages is primarily driven by efferent signals.

### Afterimage movement in adapted saccades

A plethora of evidence suggests that saccadic adaptation (i.e., the adaptive change of saccade parameters when facing systematic landing errors) affects target-localization judgments around saccades (for a nonexhaustive list, see Awater, Burr, Lappe, Morrone, & Goldberg, 2005; Bahcall & Kowler, 1999; Collins, Doré-Mazars, & Lappe, 2007; Collins, Rolfs, Deubel, & Cavanagh, 2009; Masselink & Lappe, 2021; Moidell & Bedell, 1988), which raises the question whether the same applies to the localization of afterimages. Hence, in Session 3 we compared afterimage localization in two experimental blocks that differed only in saccade trials (Figure 5a; for further details, see Session 3: Saccadic adaptation): Whereas saccade trials in the baseline block were identical to those administered in previous sessions, saccade trials in the adaptation block featured an intra-saccadic inward displacement of 2 dva. This caused primary-saccade amplitudes to decrease from an average of 13.47 dva (*SD* = 1.34) to 12.08 dva (*SD* = 1.45; *F* (1, 11) = 47.15, *η*^2^ = 0.21, *p* < .001; Figure 5b). We next tested whether the extent of afterimage movement was predicted by the magnitude of saccadic adaptation (Figure 5c): Indeed, linear mixed-effects models suggested a tight relationship between the respective changes in afterimage PNL and eye position (intercept: *β* = −0.23, *t* = −0.74, 95% CI [-0.85, 0.38]; slope: *β* = 0.85, *t* = 3.73, 95% CI [0.40, 1.31]). Note that, due to the width of distributions shown in Figure 5b, we discretized eye positions in three bins for each observer and experiment block to more accurately estimate this effect. Yet, the amount of adaptation did not seem to vary across bins, neither for eye position (*F* (2, 22) = 0.31, *η*^2^ *<* 0.01, *p* = .734, *p_GG_* = .606) nor for afterimage localization (*F* (2, 22) = 0.65, *η*^2^ = 0.03, *p* = .528, *p_GG_* = .468). These results suggest that if perceptual afterimage localization is driven by efference-copy signals, then these signals must also contain information about the adaptation of the motor command.

**Figure 5.**
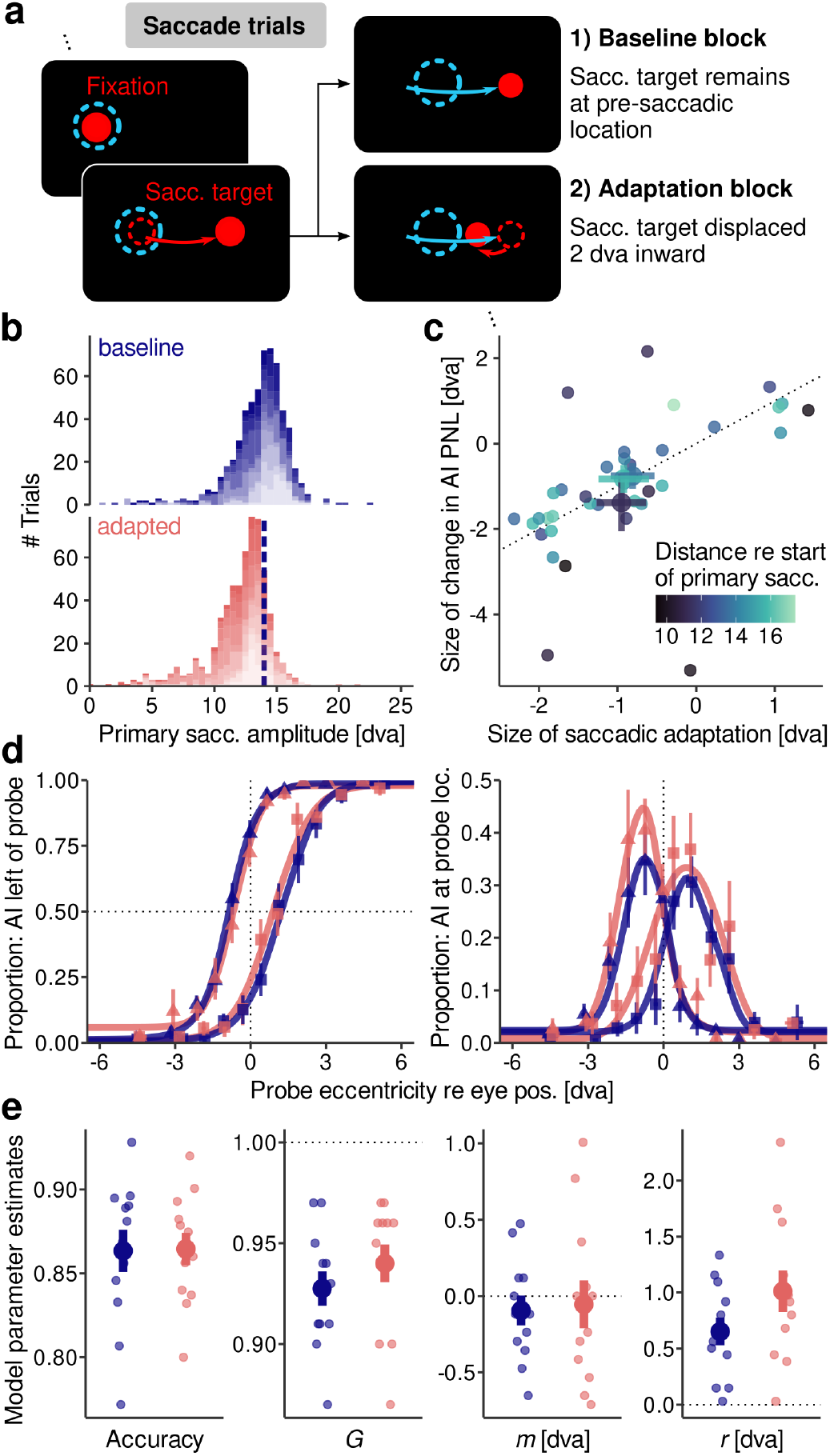
Afterimage (AI) position in adapted saccades (Session 3). **a** Illustration of inward adaptation manipulation, which was administered in two separate blocks (first baseline, then adaptation) and affected only saccade trials (for details, see Session 3: Saccadic adaptation). **b** Distributions of primary saccade amplitudes in baseline (top) and adaptation (bottom) blocks, where alpha levels indicate different observers. Dashed vertical line indicates the median amplitude in the baseline condition. **c** Size of afterimage-PNL change as a function of the corresponding change in eye position. Colors indicate average eye-movement amplitude in three eye-position bins per observer. Thick dots represent each bin’s group average (± SEM). **d** Group-level psychometric functions, logistic (left) and Gaussian (right), approximating localization judgments in baseline and adaptation block. Triangles and squares show aggregates for rightward and leftward primary saccades, respectively. **e** Classification accuracy and best-fitting model parameters for each observer and block.

To further investigate the less trivial question whether saccadic adaptation also affects the transfer function that translates efferent information to the visual domain, we once again analyzed afterimage-localization judgments relative to concurrent eye position. Interestingly, afterimage PNLs of adapted eye movements showed a reduced offset from zero (rightward: *M* = +0.20 dva, *SD* = 0.52; leftward: *M* = −0.30 dva, *SD* = 0.52; Figure 5d), but this shift remained insignificant regardless of whether it was estimated by logistic functions (*F* (1, 11) = 4.39, *η*^2^ = 0.04, *p* = .061) or Gaussian functions (*F* (1, 11) = 0.57, *η*^2^ *<* 0.01, *p* = .466). As expected from these results, estimated gains *G* slightly but insignificantly increased due to adaptation (baseline: *M* = 0.93, *SD* = 0.03; adapted: *M* = 0.94, *SD* = 0.03; *F* (1, 11) = 3.11, *η*^2^ = 0.04, *p* = .105; Figure 5e), while the center variable *m* remained largely unaltered (*F* (1, 11) = 0.29, *η*^2^ *<* 0.01, *p* = .595). The range parameter *r* increased during adaptation (*F* (1, 11) = 6.13, *η*^2^ = 0.11, *p* = .031), reflecting the larger proportion of ‘directly at’ responses with adapted saccades (Figure 5d, right). Note that this tendency, however it was generated, had no effect on afterimage PNLs. In sum, these results suggest that the visual shift of afterimages was a function of the eye movement’s size – again with gains highly similar to those measured in previous sessions – regardless of whether it was adapted or not. This finding appears different from, but (as we will argue below) is compatible with the adaptation-related PNL shifts revealed by experiments using localization relative to pre-saccadic visual targets (e.g., Bahcall & Kowler, 1999; Collins et al., 2009).

### An inverse relationship between saccade and afterimage-movement gain

Analyses up to this point have revealed two major findings. First, afterimage movement is a function of eye movement, but the process that translates motor information into the visual domain to shift afterimages operates with gains significantly below 1, thus underestimating the extent of the eye movement. Second, this hypometric gain not only explained afterimage PNLs across saccade directions and amplitudes, but also remained unaffected by the presence of post-saccadic visual references and saccadic adaptation, suggesting that it is directly related to by the motor command itself. Our finding that gains across sessions amounted to averages of around 0.94 (Session 1: *M* = 0.94, *SD* = 0.02; Session 2, no reference: *M* = 0.94, *SD* = 0.03; Session 3, baseline: *M* = 0.93, *SD* = 0.03) is reminiscent of the finding that saccades routinely undershoot their targets (e.g., Becker, 1989; Henson, 1979). We thus investigated whether and, if so, how observers’ estimated gains were related to their saccade metrics. To avoid any potential confounds with the already established relationship between eye and afterimage movement, we strictly used saccade trials (as opposed to localization trials) to estimate eye-movement metrics. We first computed primary saccade gain, that is, the ratio between actual and instructed movement amplitude, which amounted to an average of 0.976 (*SD* = 0.062), but discovered that this was likely an insufficient metric to relate to, due to the high frequency of secondary, or corrective saccades. For instance, one observer made primary saccade with gains of only 0.8 (Figure B3a), but compensated these undershoots with short-latency secondary saccades in more than 80% of trials, reaching the target with a combined gain of close to 1 (Figure B3b-c). We thus computed saccade gain as the weighted average of primary saccade gain and, if secondary saccades were made, combined gain.

For Session 1, regressions revealed a significant negative relationship between saccade gain and model-estimated afterimage gain across observers (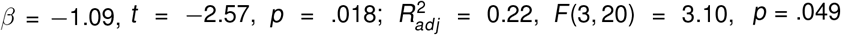; Figure 6a), irrespective of saccade direction (intercept: *β* = 0.01, *t* = 0.51, *p* = .615; slope: *β* = −0.18, *t* = −0.43, *p* = .674), suggesting that *G* parameters were lower in observers whose saccades exhibited higher saccade gain. This somewhat counterintuitive finding was replicated with data of Session 3 (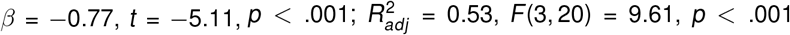; Figure 6b). Note that, despite a clear reduction of saccade gain due to adaptation that manifested itself in a change of intercept (*β* = 0.05, *t* = 4.19, *p* < .001), the estimated slope did not change significantly (*β* = 0.10, *t* = 0.69, *p* = .497), suggested relationship persisted even when saccades were adapted. To complete the picture, we also looked at the stability of estimated *G* parameters between Session 1 and 3, which were separated by a few days. Estimates were clearly positively but rather loosely related (*β* = 0.63, *t* = 2.60, *p* = .017; 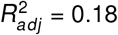, *F* (3, 20) = 2.68, *p* = .074; Figure 6c) and this relationship was largely unaffected by saccadic adaptation (intercept: *β* = −0.07, *t* = −0.33, *p* = .745; slope: *β* = 0.07, *t* = 0.03, *p* = .765). Expressed differently, the correlation between *G* parameters from the two sessions amounted to 0.49 (*t* (22) = 2.64, *p* = .015), representing a moderate relationship at best. In comparison, however, correlations of corresponding saccade gains also (only) amounted to 0.43 (*t* (22) = 2.20, *p* = .038), which hints at the possibility that both gains may be adapted flexibly over time while maintaining a highly similar negative relationship with each other.

**Figure 6.**
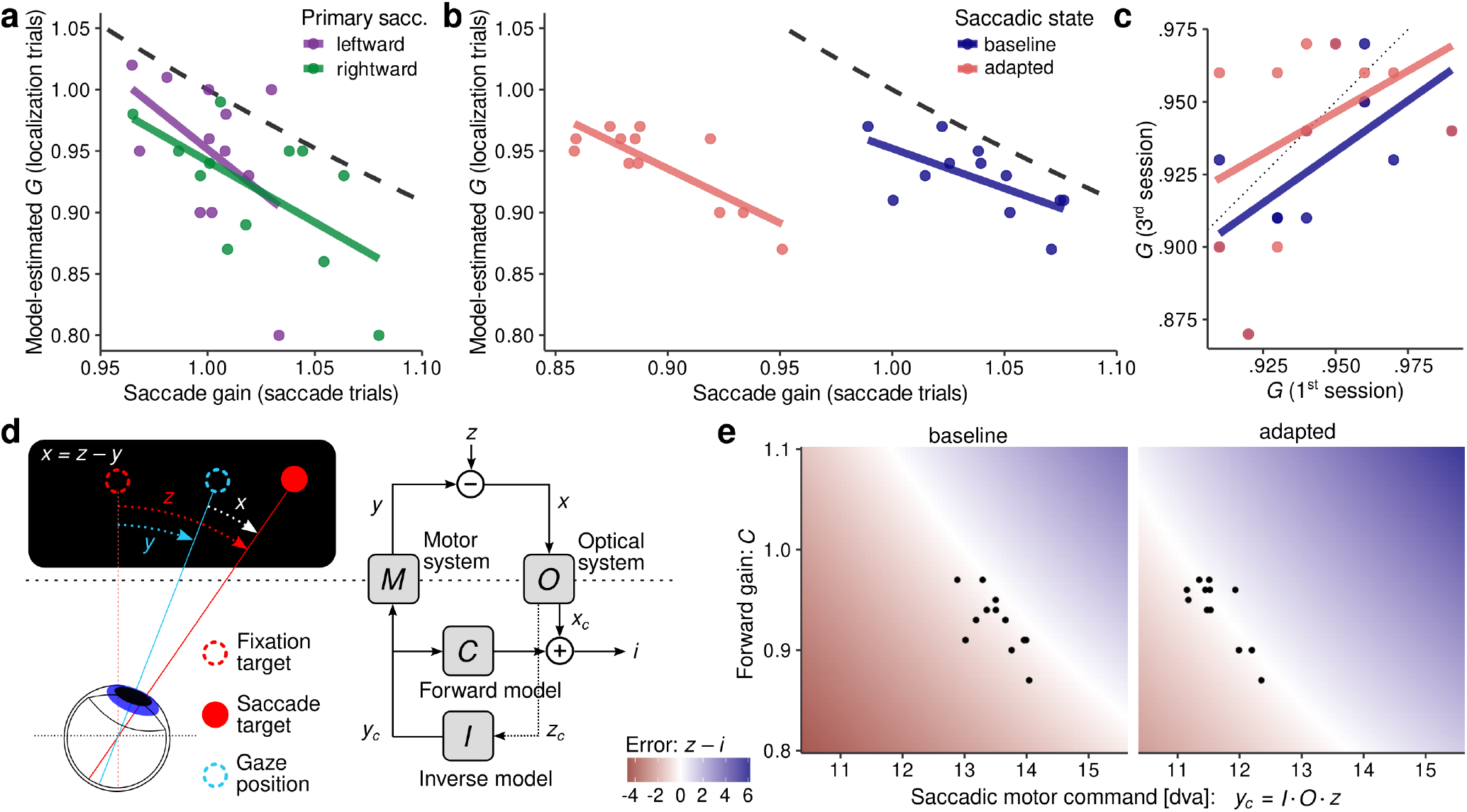
The relationship between saccade gain and model-estimated afterimage-movement gain. **a-b** Individual observers’ G parameters, estimated from afterimage-localization trials by the Gain Model, as a function of saccade gain, estimated from the same observers’ saccade trials, separately for primary saccade direction (panel a; Session 1) and experimental block (panel b; Session 3). **c** Relationship between G parameters estimated in Session 1 and Session 3, for baseline and adapted saccades, respectively. **d** Information processing structure (adapted from Mittelstaedt, 1990,, see main text and Modeling of the inverse relationship for details) as a model for head-centered visual localization, where z is the original eccentricity of the target and i is the representation of the target location. **e** Parameter space of forward and inverse model gains C and I when expressed as a function of the localization error z − i in the baseline (left, z = 13) and adapted (right, z = 11) blocks, and visual gain O set to 1.

How could this negative relationship between *G* and saccade gain be explained? For a tentative explanation in terms of control theory, we adapted an information processing structure proposed by Mittelstaedt (1990). This simplified system (Figure 6d; see Modeling of the inverse relationship for a detailed description) is equipped with a motor system, that transforms a motor command *y_c_* into a movement *y*, and an optical system that transforms physical eccentricities *z* or *x* into coordinates in visual space. To localize objects in space, it combines the efference copy of the motor command with a visual coordinate *x_c_* . Critically, the efference-copy signal (e.g., the number of steps with a stepper motor) must be translated to a visual signal (e.g., the number of pixels on a camera image) to be of any functional use for visual localization – which is the role of the forward model with gain *C* (Equation 7). This visuomotor gain *C* would be the analogue of the parameter *G* in our Gain Model: It translates a quantity observed in the motor domain (i.e., eye-movement amplitude) into a visual quantity (i.e., amplitude of afterimage movement) and therefore forms the basis for predicting the visual consequences of saccades. But how is the motor command *y_c_* determined? Masselink and Lappe (2021) proposed an inverse model (here with inverse gain *I*) that transforms the visual representation of the pre-saccadic target eccentricity *z* into a motor command that can reach that target (Equation 9), allowing target localization based on the predicted visual consequences of the motor command and the (post-saccadic) target eccentricity (Equation 10). Because in the case of afterimage localization there is no physical target to localize, the perceived location of the afterimage reduces to Equation 11:

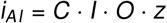

This definition means that the movement of afterimages depends on the actual eye movement, which ultimately depends on motor gain *M*, only in so far that they share the same motor command: What is copied is the “effort of will” (Willensanstrengung; von Helmholtz, 1867) – an assumption plausibly supported by the observation of subjective movement of the visual environment when saccades were attempted but unsuccessful due to extraocular paralysis (Stevens et al., 1976). It also becomes evident that, in order to achieve an efference-based spatial localization of afterimages that approximates the initial saccade target eccentricity (i.e., minimizing the error *z* − *i*), the inverse gain *I* (and thus the resulting motor command *y_c_*) and forward gain *C* should ideally be inversely related. The negative relationship between saccade gain and estimated *G* described above clearly followed this model prediction, irrespective of whether saccades were adapted or not (Figure 6e). While the small increase of *G* in the adapted block was not significant and also Bayesian ttests (Rouder, Speckman, Sun, Morey, & Iverson, 2009) did not provide evidence for any change of *G* in the adaptation block (*BF* = 0.951), these results and modeling suggest that, in order to match the intra-saccadic displacement introduced in saccadic-adaptation paradigms, it may be sufficient to adjust the size of the motor command via the inverse model, as – thanks to the consistent (false) post-saccadic visual feedback – the contingency between motor command and visual consequence *C* remains unaltered. Note that *G* below 1 cause localization to sys-tematically fall short of veridicality and that the impression of veridical localization in the adapted case is a false one, as the adaptation of eye movements did not reach the size of the intra-saccadic target displacement (i.e., 2 dva). Yet, grounding empirical results in this simple model strengthens the notion that phenomenal movement of afterimages across saccades reflects the outcome of predictive processes that estimate the visual consequences of saccadic motor commands. These purely efference-based predictive processes might not be veridical, but they may not need to be, because post-saccadic visual feedback is available during natural viewing to complement visual localization. Our results suggest that, even though they are constrained by an inverse relationship, the calibration of inverse and forward models individual for each observer and may be learned through repeated exposure to the consequence of one’s own saccades.

## Discussion

Before dedicated eye-tracking apparatuses were developed, observing the movement of afterimages in egocentric visual space was the primary technique to assess one’s own gaze position (Wade, 2015). Up to this day, however, very little is known about the extent of the – phenomenologically extremely convincing – congruency between afterimage position and eye position. In consideration of the broader questions of oculomotor control and visual stability, this question is highly interesting, as afterimage movement across saccades must be a consequence of the attempts of the system to account for the visual consequences of its own movement. In fact, subjectively, there is a clear dissociation between the position of an afterimage on the one hand and the position of the visual world on the other: The afterimage changes across saccades while remaining static in the retinotopic frame of reference; conversely, the world is stable across saccades while moving in the retinotopic frame of reference. During unusual viewing conditions, for instance, when pressing the eye ball (Ilg, Bridgeman, & Hoffmann, 1989) or performing voluntary oscillopsia (Enright, 1994), the visual world moves, but afterimages remain steady. We thus hypothesized that observing afterimage movement across saccades could provide new insights into the system’s internal representation of saccade-induced visual consequences. In this study, we systematically measured the perceived position of afterimages induced prior to a primary saccade, while continuously tracking observers’ true eye positions. To express the two variables as a function of each other, we fitted a tracking model to the data that assumed a set linear relationship between changes in eye and afterimage position, respectively, that was characterized by the gain parameter *G*. Performing experiments in the well-controlled environment of total darkness assured that afterimage position was not contaminated by visual references. That way, the estimated gain could be interpreted as the system’s estimate of the size of its own eye movement relative to the size of the eye movement actually made. Our results can be summarized by four major findings, discussed below.

First, average afterimage-movement gains amounted to 0.94, confirming a tight, but hypometric relationship between eye movement and resulting afterimage movement. It should be noted that gains of 1 would occur either if the system had perfect knowledge about the exact size of the executed eye movement, or if it had no such knowledge at all and would perform localization based purely on comparing the retinotopic locations of afterimage and probe. That gains were significantly and consistently below 1 rules out either of these alternative hypotheses and shows that, despite their tight relationship, afterimage position did not equate to eye position. Note that this result does not contradict the assumption that afterimages remain fixed in retinotopic space: If localization is performed in egocentric space, then retinotopic information must be combined with eye-position information, which (in the model assumed here) is based on an efference-based and feedforward motor-to-visual transformation estimating the visual shift induced by a given saccadic motor command. If this transformation is hypometric, so will be the apparent movement of any visual image fixed in retinotopic coordinates. Interestingly, despite their hypometry, our gain estimates were considerably closer to 1 than those reported by other investigators. For instance, Grüsser et al. (1987) found average gains of 0.5 when making saccades at a rate of 1.5 Hz and gains of virtually 0 at saccade rates approaching 4 Hz, reproducing the phenomenon of static afterimages during voluntary oscillopsia (Enright, 1994). Notably, they also reported higher gains when saccade rates were further reduced (e.g., 0.72 at 0.2 Hz; Grüsser et al., 1987, their Fig. 4a), suggesting the possibility that in single-saccade paradigms gains may well be even higher, as it was the case in our study. A major difference to this previous study, however, was that afterimage-localization trials were intermixed with saccade trials, in which saccade targets remained continuously present as visual references across eye movements. Thus, despite the otherwise reference-free visual environment, there was ample opportunity for the system to calibrate its motor commands – that is, to adjust the gains of every subsystem (Figure 6d) – with preand post-saccadic visual feedback. In this perspective, it may not be surprising to find afterimage-movement gains close to 1, as thanks to repeated observations, accurate predictions regarding the sensory consequences of motor commands could be made. In the assumed model, the forward gain *C* governs this motor-to-visual transformation. Mittelstaedt (1990, 1994) provided formal proof (see Equation 8) that localization would be independent of the actual size of the motor command, if *C* = *O · M*, that is, if the forward gain knew and incorporated visual and motor gains. Indeed, that post-saccadic localization can be independent of oculomotor error has been shown previously (Collins et al., 2009). From our data, we cannot estimate these variables, as dedicated localization tasks without saccades would be needed to estimate visual gain (e.g., Masselink & Lappe, 2021). Previous evidence, however, suggests an overestimation of visual target eccentricity without saccades, but an underestimation when saccades were made to localization targets (Morgan, 1978). Furthermore, when measuring saccade gain by assessing motor output (*y*), it is unclear how much of the result should be attributed to motor or inverse gains, respectively. In other words, if a saccade lands short of its target, was the saccadic motor command not properly executed by the extraocular muscles (*M* too small) or was the motor command unfit to reach the target to begin with (*I* too small), or both? For this case, measuring the afterimage movement could provide an interesting venue, as its size should in principle depend on the motor command (*y_c_*), not actual motor output (*y*). The forward gain *C*, and our fitted parameter *G*, may thus reflect the system’s experience with the typical sensory consequences of motor commands issued in natural environments. Given the proneness of saccades to undershoot their targets by 5-15% (for a review, see Becker, 1989) and assuming that this error is related to (possibly to some extent hardwired) visual and motor gains, then it would only be reasonable of the system to predictively underestimate the visual consequence of any given motor command. Interestingly, Masselink and Lappe (2021), who had also found hypometric forward gains in their experiments, championed a similar argument but with its direction being opposite, that is, that “systematic inaccuracies of saccades stem from inaccuracy of internal movement representations” (p. 24).

Second, how does this perspective fit with the notion that forward and inverse gains should have an inverse relationship? We confirmed this notion by establishing negative linear relationships between saccade gain and afterimage-movement gain that were present consistently across saccade directions and experimental sessions. As pointed out above, saccade gain is the final result of visual, inverse, and motor gains, so for the simulations of localization error (Figure 6e) we set visual and motor gains to 1. The inverse relationship is somewhat unintuitive, as we have seen that afterimage movement scales positively and linearly with the size of the eye movement and therefore with the driving motor command, which itself depends on the inverse gain. Using the example of a camera mounted on a stepper motor, the inverse relationship can be understood. To determine the number of steps needed to foveate a target at some eccentricity measured in pixels (i.e., the inverse model), the system would need to know how many steps would lead to a shift of one pixel (the inverse gain). Conversely, to make a prediction about the amount of pixel change that would result from a certain stepper activation (i.e., the forward model), the system would need to know the shift in pixels resulting from one step of the motor (the forward gain). From this example, it is immediately evident that these two gains must be inversely related, as they both refer to the same contingency between camera and motor. Importantly, the sensorimotor system is likely not perfect: Slack in the stepper motor or optical distortions in the lens of the camera could complicate the establishment of a straightforward rule, which is why these gains must remain plastic, for instance to compensate for unexpected errors (e.g., in saccadic adaptation) or gradually introduced errors (e.g., due to fatigue of the extraocular muscles). In this sense, when considered on the individual-observer level, the relationship can be interpreted as an adjustment to the inherent visual and motor gains of the system. On the one hand, as demonstrated by Mittelstaedt (1990), the forward gain should scale with visual and motor gains. For example, motor gains below 1 would cause saccades to consistently fall short of their targets, a tendency that would be incorporated in the prediction of visual consequences by reducing the forward gain. On the other hand, in the face of repeated prediction errors the system will adjust its motor commands to correct for them. With motor gains below 1, inverse gains would be increased to up-regulate motor output and thereby compensate for shortcomings of the motor system. As the example shows, adjustment of inverse and forward gains would again be made in opposite directions, as previous saccadic-adaptation experiments have already shown (Masselink & Lappe, 2021), leading to a calibration of sensorimotor predictions that is tailored to the properties of each individual system.

Third, we found that saccadic adaptation did not alter the tight link between eye and afterimage movement: Whether adapted or not, the size of afterimage movement scaled proportionally with the size of the eye movement – with largely the same gain. This result shows that the efference copy carries information about adapted motor command for visual localization, thus providing strong evidence against the opposite hypothesis proposed to explain the shift of PNLs of saccade targets due to saccadic adaptation (Collins et al., 2009). When subjected to a saccadic-adaptation paradigm, in which the saccade target is consistently stepped inwards, it seems to be sufficient for the system to reduce its inverse gain to downregulate motor input, thereby shortening saccade amplitude and minimizing visual error. As this down-regulated motor input is then passed to the forward model, the latter will automatically – that is, even without having to change its gain – predict the smaller visual change that would result from the movement. Ironically, this prediction will even match post-saccadic visual feedback, due to the consistency of the experimentally induced intra-saccadic displacement. If in the adapted state the intra-saccadic displacement were to be suspended and the saccade target remained in its original pre-saccadic position, then unexpected visual offset (*x_c_*, see Equation 7) would be introduced, making the target appear as if having moved in an outward direction. Thus, constant forward gain would be compatible with findings of adaptation-induced PNL shifts (Bahcall & Kowler, 1999; Collins et al., 2009). Whereas we found no change in afterimage-movement gain due to adaptation, Masselink and Lappe (2021) found small but significant increases in their corollary-discharge gains (i.e., their equivalent of the gain *C* proposed by Mittelstaedt, 1990) as motor gains (i.e., their equivalent of the inverse gain *I*) decreased due to inward adaptation of saccades. The source of these differences is unclear, but most likely lies in the experimental paradigm (their observers localizing pre-saccadic flashes after primary saccade offset instead of afterimages) and study design (their sample size being larger than ours). It is thus not impossible that our study’s sample size did not allow for statistically securing very small increases in forward gain. Alternatively, it may well be possible that sensorimotor learning had not reached its asymptote yet, or that the forward model was adjusted more sluggishly than, or lagging behind, the inverse model.

Fourth and finally, our results showed that perceived afterimage position was independent of post-saccadic visual feedback, providing further support that phenomenological movement of afterimages was indeed a consequence of efference-based prediction of visual consequences. To elaborate, based on the contingency illustrated in Figure 6d, the continuous presence of the saccade target allows for the direction estimation of the saccade’s visual consequences by measuring the post-saccadic offset *x*, provided the assumption of stability of visual objects in space (Born, 2019; Deubel, Schneider, & Bridgeman, 1996). In that case, if the actual extent of the visual change were known and taken into account then one might expect to see a stronger alignment of eye and afterimage movements, that is, a higher afterimage-movement gain, in comparison to the total-darkness situation where the system could only rely on the accuracy of extra-retinal sources of information or its sensorimotor predictions. Similar results could be expected if displacement of afterimages was driven by combining extra-retinal and retinal feedback (Bridgeman et al., 1994), or by some sort of postdictive process that estimated the visual consequence of each saccade based on the resulting visual error with reference to the saccade target. This is especially true in the false-feedback condition where post-saccadic visual errors would trick the system into believing that the executed saccade overshot the target. That afterimage-localization judgments, however, remained unaffected by such manipulations suggests that these factors were likely not taken into account. It could be argued that the duration of 150 ms, for which saccade targets remained visible after saccade onset (see Figure 4a), may have been too short to reliably act as post-saccadic reference. While this short duration was chosen deliberately to avoid potential interactions with the emergence of afterimages, previous findings suggest that such post-saccadic durations are sufficient to induce saccadic suppression of image displacement (Deubel et al., 1996), mask motion signals (Duyck, Wexler, Castet, & Collins, 2018), and provide reliable visual cues for motion amplitude (Rolfs, Schweitzer, Castet, Watson, & Ohl, 2025). In addition, even shorter target durations were capable of inducing saccadic adaptation (Panouillères, Gaveau, Socasau, Urquizar, & Pélisson, 2013), suggesting that even brief, early post-saccadic visual references could be used to estimate saccadic error. Our results thus strengthen the model’s assumption that, if the afterimage movement is truly a product of the forward modeling of saccadic consequences, then it should depend exclusively on the efference copy of the motor command, and not on the actual saccade size.

To conclude, psychophysical localization of afterimage position across saccades revealed that the size of afterimage movement directly scaled with the corresponding size of eye movement. Importantly, estimated gains were high but consistently hypometric, confirming previous measurements made with target-localization paradigms (Masselink & Lappe, 2021). As afterimage-movement gains scaled inversely with saccade gain and remained unaffected by not only post-saccadic visual references but also saccadic adaptation, there are good reasons to assume that afterimages move based on feedforward predictions of the visual consequences of saccades, which is in parsimonious accordance with a basic solution for head-centered visual localization based on efference copy of the motor command (Mittelstaedt, 1990). In contrast, it is rather implausible that a solution based on proprioceptive eye-position signals could also give rise to this localization, as we could confirm the observation made by Grüsser et al. (1987) that afterimage movement was limited to active eye movements and could not be induced by passively moving the eye, for instance, by pressing the eye ball or tapping the outer canthus. The technique of tracking afterimages across eye movements thus enables direct and efficient insights into the motor-to-visual transformations that underlie space constancy, and that could so far only be estimated indirectly, that is, by comparing pre-saccadic with post-saccadic localization judgments (cf. Masselink & Lappe, 2021). In this study, we investigated afterimage movement with the system being in a calibrated state, achieved by presenting trials with static and continuously present saccade targets. If such trials had not been introduced, it is unclear whether our results would hold. Moreover, while we only assessed the spatial characteristics of afterimage movement, another critical question to be addressed by future studies would be the time course and temporal precision of the underlying motorto-visual forward model. That Grüsser et al. (1987) estimated afterimage-movement deadtimes of 180-330 ms, suggesting considerable sluggishness given the brevity of fixations, raises the question whether afterimage movement is even representative of the assumed feedforward predictive process. Revisiting these results – which, notably, also yielded substantially lower gains than we reported here – with our paradigm may thus provide insights in to what extent the mechanisms that underlie afterimage movement could actually contribute to visual stability, on both spatial and temporal scales. In this sense, our study is only a starting point, demonstrating the great potential of the afterimage-tracking technique for investigating the sensorimotor computations routinely performed during active vision.

## Methods

### Participants

Twelve observers (five female, age range [22, 46]), two of them authors of this study, participated in the experiment. All were instructed about the task procedure, informed about the purpose of the experiment, and gave written informed consent prior to their participation in the study. All had normal or corrected-to-normal vision (one person wore glasses), as determined by the Snellen test. Seven participants had right ocular dominance (established by a variant of the Porta test) and ten were righthanded. The experiment was conducted in agreement with the latest version of the Declaration of Helsinki (2013) and was approved beforehand by the Ethics board of the Department of Psychology at Humboldt-Universität zu Berlin.

### Apparatus

Experiments took place in a completely dark room devoid of any systematic light sources. The experimenter was seated in a separate room, communicating with participants via intercom. Eye tracking of a participant’s dominant eye was performed using an Eyelink 1000 Plus desktop mount system (SR Research, Osgoode, ON, Canada) using a sampling frequency of 1000 Hz with standard link and file filter settings. To render the position of the eye tracker indiscernible in low-light viewing conditions, we applied a 940 nm infrared illuminator with an additional diffusion filter fitted on top. Interfacing between eye tracker and experimental control was realized using PsychoPy version 2022.2.5 (Peirce, 2007) and the dedicated pylink released as part of the Eyelink developers kit provided by SR Research.

To avoid visual references typically created by the backlight of standard stimulus-presentation monitors or the beams of projection systems, we utilized a custom-built LED screen with a resolution of 48 × 32 RGB LED pixels. Specifically, the screen consisted of six 16×16 DotStar matrices (Adafruit Industries, New York City, NY, USA) which were driven–connected via six 74AHCT125 level shifters (for a related setup, see Schweitzer, Watson, Watson, & Rolfs, 2019)–by an Arduino Due microcontroller hosting the serial interface to the stimulus computer. The dimensions of the screen amounted to approximately 48.6 × 32.4 cm and participants were seated at a distance that allowed the visual angle between adjacent LED pixels to amount to one degree of visual angle (dva). An acrylic glass diffusor was fitted on top of the LED panels so that diode electronics remained hidden and illuminated pixels appeared like Gaussian blobs.

Despite its comparably low spatial resolution, our custom-built screen had the advantage of high temporal resolution, allowing single LED activations as short as 3.5 ms on average (Figure B1a). To log information about the precise onset times of LEDs, TTL triggers were sent by the microcontroller to the eye tracker as soon as LEDs were activated. The screen achieved short display latencies, that is, the time from sending the serial command to the physical activation of the LED, which amounted to an average of 4.3 ms (SD = 1.4). In the tested sample of 28163 saccade-triggered presentations, we found a minimum and maximum latency of 0 ms and 10 ms, respectively, and used these values as conservative estimates of display latency for trial exclusions (see Preprocessing). We experimentally confirmed that TTL triggers and LED activity were temporally aligned with zero lag (Figure B1b) and measured the luminance and color of our stimuli, as well as the linear gamma of the utilized LEDs (Figure B1c).

To provide manual responses, participants used the LeftArrow, RightArrow, DownArrow, and Space keys on a standard US-layout keyboard. Sounds were played with a Sony SRS-XB12 speaker.

### Stimuli

All visual stimuli were dots produced by activating single LEDs placed behind the diffusor screen. Due to the resulting diffusion, they appeared similar to Gaussian blobs with a standard deviation of 0.25 dva. Fixation and saccade targets were of minimum-luminance red color (RGB: [1,0,0], CIE x/y coordinates: 0.68/0.31, 0.4 cd/m^2^) and afterimage inducer pulses were of maximum-luminance white (RGB: [255,255,255], CIE x/y: 0.31/0.27, 230.1 cd/m^2^). Inducer pulses were always shown as a sequences of twelve presentations with a duration of 50 ms and an inter-stimulus-interval (ISI) of 10 ms in between. As the auditory response cue the ‘qbeep’ file was used, which is provided by SR Research along with the Eyelink calibration routines for PsychoPy.

### Procedure

The experiment consisted of three sessions – each using a different experimental manipulation – performed on separate days, each with an uninterrupted duration of approximately one hour. In order to achieve a sufficient level of dark adaptation, a pre-experiment (typically 10-20 minutes) unrelated to the localization of afterimages was performed at the beginning of each session and then directly succeeded by the afterimage task. Regardless of condition or block, a break was introduced after every 110 trials, in order to allow participants to rest and to re-run the calibration of the eye tracker.

Generally, in all types of trials, participants were instructed to make leftward and rightward horizontal saccades in response to the presentation of a peripheral saccade target. These visually guided saccades triggered stimulus manipulations and will henceforth be referred to as primary saccades. Fixation positions displayed in the beginning of each trial were randomly varied horizontally and vertically by up to +/3 dva. To facilitate the fluency of the task, primary saccades were always made in alternating direction, unless a trial had to be repeated at the end of each experimental block. Such repetitions were automatically triggered if participants failed to pass fixation control, made no saccade up until five seconds after the presentation of the saccade cue, or made a saccade before the saccade cue was presented. In addition, to make sure that adequate saccades were made, trials were repeated in which the primary saccade amplitude was less than 60% of the instructed saccade amplitude. Online saccade detection, in order to trigger potential stimulus manipulations, was performed using the algorithm described by Schweitzer and Rolfs (2020) with parameters *λ* = 10 and *k* = 3, and without applying a direction criterion. Across the group of participants, average online saccade detection occurred on average 8.1 ms (SD = 0.25) after physical (i.e., offline-detected) saccade onset, with individual latency distributions having an average standard deviation of 2.4 (SD = 0.53). Saccade offsets were detected with the same algorithm, but–to achieve a more strict criterion for saccade offsets than onsets–eight below-threshold samples were set as a criterion. All trials, regardless of condition, started with fixation control, where participants were required to maintain fixation within a circular boundary with a radius of 2 dva around the fixation target for a randomly sampled duration ranging from 400 to 600 ms. From the onset of the first fixation within the boundary, five seconds were given to pass fixation control, else the trial was aborted and a re-calibration of the eye tracker was triggered.

#### Pre-experiment

Pre-experiments consisted of 304 trials, of which 160 were saccade trials. In saccade trials, participants made single primary saccades with an instructed amplitude of 13 dva to follow the horizontal displacement of the fixation target, provided that fixation control was passed. In the remaining 144 test trials, which were signaled by a beep sound presented 200 ms after the online detection of the primary saccade, participants reported the direction of the intra-saccadic displacement of the saccade target by pressing the LeftArrow key for leftward displacements, the RightArrow key for rightward displacements, and the DownArrow key to indicate that the target remained stationary. Specifically, upon saccade-onset detection, the saccade target could be displaced horizontally by −4 to 4 dva (with a step size of 1 dva) with respect to its presaccadic position. Critically, there were four presentation conditions in which these displacements could be shown (Figure A1a-b). First, there was the saccadic-suppression-of-displacement (SSD; Bridgeman, Hendry, & Stark, 1975) condition, in which the jump of the saccade target from one location to another was without temporal delay and both pre- and post-saccadic saccade targets were continuously present until the end of the trial. Note that all saccade trials belonged to this condition. Second, in the presaccadic blanking condition the pre-saccadic saccade target was flashed for 50 ms, while the post-saccadic target was presented without delay and remained continuously present until the manual response was given. Third, in the post-saccadic blanking condition the pre-saccadic saccade target remained continuously on until saccade detection and then extinguished for 200 ms (i.e., the blanking duration; Deubel et al., 1996) until the (possibly displaced) post-saccadic target appeared together with the auditory response cue. Finally, in the pre- and post-saccadic blanking condition the pre-saccadic saccade target was flashed for 50 ms and the post-saccadic saccade target appeared after the blanking duration. Taking into account each participant’s individual saccade-latency distribution, their average pre-saccadic blanking amounted to 90.2 ms (SD = 25.9). Considering the latency of the saccade-triggered ISS and individual saccade durations, the post-saccadic blanking interval ended 170.7 ms (SD = 2.0) after saccade offset.

Two trial repetitions per experimental condition (saccade direction × ISS size × presentation condition) were run per session, totalling six data points per experimental cell across all three sessions.

#### Main experiment

Whereas pre-experiments dealt with the localization of targets displaced across saccades, main experiments dealt with the localization of retinal afterimages relative to spatial cues presented after saccades were executed. As in the pre-experiment, participants were performed both saccade and test trials. Again, trials in the main experiment were initiated with the presentation of a single fixation dot and subsequent fixation control. Once passed, in saccade trials the saccade target was displayed, whereas in test trials a bright inducer pulse (see Stimuli) was presented which participants were instructed to continue to fixate. Following a randomly sampled delay ranging from 75 to 150 ms, the saccade target was presented. In sessions 1 and 3, the saccade target was flashed for 50 ms, whereas in session 2 the saccade target was extinguished once the saccade was detected. After the saccade was executed, a beep sound informed participants that they should press the Space key as soon as they had a clear view of the pre-saccadically induced afterimage. As the subjective appearance of the afterimage may vary across participants and throughout the course of the experiment, this ready-response was fully self-paced. Participants average ready-response latencies amounted to a mean of 1165.1 ms (SD = 455.2) after primary saccade offset, that is, on average 1476.4 ms (SD = 469.9) after inducer offset, making it unlikely that afterimages could have directly affected the execution of the primary saccade. Once participants reported the presence of the afterimage, a sequence of up to seven probes was flashed, their horizontal spatial positions randomly selected from −3 to +3 dva relative to the (by now long extinguished) pre-saccadic saccade target. These probes were of the same color as fixation and saccade targets, but were presented for the shortest possible duration (i.e., on average 3.5 ms; see Apparatus) to prevent probe-induced afterimages or spatial interactions with the induced afterimage, and to discourage their use as visual references. After each appearance of a single probe, participants indicated whether the position of the afterimage was located right of (RightArrow key), left of (LeftArrow key), or directly at (DownArrow key) the probe location. Participants could either continue to respond until all probe locations were shown or could opt out by pressing once more the Space key, thereby indicating that a clear afterimage was no longer present or discernible.

In each session, four test trial repetitions per experimental condition (saccade × direction probe location × experimental condition) were run. Note that, owing to the fact that in each test trial up to seven probe locations were presented, a single trial would not return a single but multiple data points, significantly increasing the statistical power resulting from the task. Specifically, each participant’s average number of responses was 6.6 (SD = 0.4) and in 80.9% (SD = 18.3) all seven responses were given. In comparison, in only 0.35% (SD = 0.38) of all trials participants reported to not have had a clear view of the afterimage after the saccade and thereby opted out of responding.

#### Session 1: Saccade amplitude

The goal of this first session was to familiarize participants with the afterimage localization task and to establish the effect of saccade amplitude on the perceived extent of afterimage motion. To investigate this question, three primary saccade amplitudes – 10, 13, and 16 dva – were instructed in both saccade and test trials. With respect to their presentation condition, saccade trials were identical to those employed in the pre-experiment. In test trials, after successful fixation control, saccade targets were flashed for 50 ms and remained extinguished for the rest of the trial, so that participants’ primary saccades landed in total darkness and, as a consequence, their localization judgements were entirely unaffected by visual references. In total, and not counting trial repetitions, 240 saccade trials and 168 test trials were scheduled in the main experiment in session 1.

#### Session 2: Visual references

In the second session, the role of post-saccadic visual references was investigated. Like in session 1, saccade amplitudes of 10, 13, and 16 dva were instructed in order to induce greater uncertainty about the impending saccade size. Unlike session 1 though, these amplitudes were selected by random choice on each trial. To manipulate the availability of post-saccadic visual references, three presentation conditions were devised. First, in the no-feedback condition, saccade targets were extinguished as soon as the saccade was detected. Second, in the feedback condition, saccade targets remained present for 150 ms after saccade detection and were then extinguished. That way, depending on saccade duration and detection latency, targets remained visible for participants for 120.4 ms (SD = 3.8) after saccade offset on average. Third, in the false-feedback condition, saccade targets were also set to remain present for 150 ms after saccade detection but at a location shifted horizontally inward by 2 dva with respect to the pre-saccadic saccade target, thus emulating a condition in which the saccade overshot the saccade target. As these shifts occurred unpredictably on a comparably small number of trials and strictly during saccades, they were not noticed (Bridgeman et al., 1975). Moreover, owing to the fact that targets were only briefly visible after primary saccades and false feedback was rare and inconsistent, as well as constantly flanked by saccade trials with unaltered visual feedback, saccadic adaptation did not take place. As in session 1, a total of 240 saccade trials and 168 test trials was scheduled in session 2.

#### Session 3: Saccadic adaptation

In the third and last session, we investigated the impact of saccadic adaptation on afterimage localization. To achieve this, we divided the main experiment of session 3 in three blocks – a baseline block, an adaptation block, and an adapted block. In the baseline block (196 trials), saccade amplitudes of 13 dva were instructed. In this block, 140 saccade trials and 56 test trials were scheduled. In the adaptation block (154 trials), amplitudes of 13 dva were instructed but saccade targets were shifted inward (i.e., left for rightward saccades and right for leftward saccades) by 2 dva once primary saccades were detected online. 140 saccade trials and a reduced number of 14 test trials were presented in this block. Finally, in the adapted block (196 trials), inward target shift were continued in 140 saccade trials to maintain saccadic adaptation (Collins et al., 2009) while 56 test trials probed afterimage localization in this adapted state. Note that test trials in all blocks were identical, regardless of saccadic adaptation: Similar to session 1, saccade targets were flashed for 50 ms at a fixed eccentricity of 13 dva and no post-saccadic visual feedback was provided. Localization probes, too, were always presented relative to the pre-saccadic saccade target to make sure that potential changes in the localization of afterimages could be attributed to saccadic adaptation and not to differences between test trials. To minimize possible awareness of changing experimental manipulations, the three blocks were seamlessly concatenated.

### Preprocessing

Each trial’s eye-tracking data, sampled at 1000 Hz, was scanned for missing data and blinks. Blink events were detected based on pupillometry noise (Hershman, Henik, & Cohen, 2018). Prior to saccade detection, gaze position data was linearly interpolated across an interval ranging from 20 samples before and after blink onset and offset, respectively. Basic saccade detection was performed with the Engbert-Kliegl algorithm (Engbert & Kliegl, 2003; Engbert & Mergenthaler, 2006), setting the velocity scaling factor to 6 and the minimum-duration criterion to 10 samples. In addition, clusters of saccade candidates separated by less than 20 samples were merged and direction-based detection of post-saccadic oscillations (PSOs; Schweitzer & Rolfs, 2022) was run for each resulting saccade interval, using a direction cut-off of 90 deg. For subsequent analyses, PSO onset, if it was detected, was defined as the time of saccade offset instead of conventional saccade offset, as the peak of the PSO constitutes a more conservative estimate of physical saccade duration (Deubel & Bridgeman, 1995a, 1995b). In order to avoid potential confounds due to PSOs, however, saccade amplitude was defined as the spatial distance traveled between saccade onset and PSO offset. Only those saccades were kept that contained no missing eye-tracking data samples.

Individual trials were excluded in case participants did not pass fixation control, executed their primary saccade prior to the presentation of the saccade target, made no primary saccade whatsoever, or reported that they had no clear post-saccadic view of the afterimage. Furthermore, trials were excluded in case online saccade detected failed or occurred with a latency of over 30 ms, as well as when saccade-triggered stimulus manipulations – taking into account minimum and maximum presentation latencies of 0 and 10 ms (see Apparatus) – did not occur strictly between offline-detected saccade onset and (PSO-defined) offset. Finally, trials were excluded in which the proportion of missing eye-tracking samples in a trial, for instance, due to blinks or insufficient tracking quality, was larger than 20% or the primary saccade could not be detected.

Following these criteria, an average of 2.7% (SD = 3.3) of trials in the pre-experiment and 13.1% (SD = 9.3) of trials in the main experiment were excluded. As some of the exclusion criteria were already already detected online, the final number of test trials amounted to 144.4 (SD = 4.2) in the pre-experiment (144 planned), and in the main experiment 167.5 (SD = 23.1) trials remained in session 1 (168 planned), 156.1 (SD = 12.0) in session 2 (168 planned), and 113.1 (SD = 14.5) in session 3 (126 planned).

### Analyses

To estimate parameters of psychometric functions individually for each observer, we used the R package ‘nlme’ (Pinheiro, Bates, DebRoy, Sarkar, & R Core Team, 2020). Fits were performed separately for experimental conditions, encompassed all observers, and allowed all function parameters (see below) to vary freely. Two types of functions were fitted to the data. First, a logistic function with the form

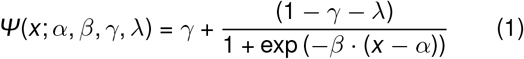

form

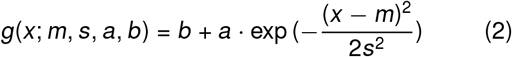

approximated the proportion of binary responses (i.e., ‘left’ vs ‘right’ or ‘inward’ vs ‘outward’). For these analyses, DownArrow key presses were transformed into binary responses by seed-controlled random sampling (García-Pérez & Peli, 2011). Second, a Gaussian function with the approximated the proportion of ‘stationary’ or ‘directly at’ responses (see Procedure). In both cases, *x* represented a measure of spatial distance in degrees of visual angle (dva), such as the size of the intra-saccadic displacement in the pre-test (Figure A1c,e) and probe eccentricity relative to start of the saccade (Figure 1c) or probe eccentricity relative to registered eye position or modeled afterimage position (Figure 2d-f) in the main experiment. To aggregate proportions in the latter cases, continuous data in each condition was discretized in 11 bins, each with an equal amount of localization judgments.

For hypothesis testing, estimated parameters were subjected to type-2 repeated-measures ANOVAs, as implemented in the ‘ez’ R package (Lawrence, 2016), with Greenhouse-Geisser corrections for sphericity (corrected p-values denoted *p_GG_*). Group-level psychometric functions were computed by averaging all observers’ individual psychometric function predictions and could therefore deviate from their original parametric shape. All error bars represent arithmetic means *±* one standard error of the mean.

### Modeling

#### Estimation of afterimage movement gain

Prior to estimating afterimage position with respect to eye position, we extracted a number of variables from individual trial data (see Figure 2a for schematics). First, we extracted the gaze positions at saccade start (*g*_0_) and landing (*g*_1_) and computed median gaze position in a 20-ms time window after each probe onset, providing a sequence of up to nine gaze positions (in the case of seven localization-probe presentations) represented in the variable *g_i_*. Note that we analyzed only the horizontal dimension of gaze positions, as this was the dimension along with observers performed afterimage localization. Second, the corresponding probe positions, denoted *p_k_*, were extracted from each trial’s data. As probes were only presented after the primary saccade was executed and observers reported the appearance of the afterimage, the first probe location *p*_1_ is matched with gaze position *g*_2_. Third, in order to control for the retinal eccentricity of the afterimage, we computed the gaze position offset from the original fixation location during the presentation of the inducer pulse and corrected subsequent gaze positions accordingly (Figure 2a, top). As shown in Figure B2a, afterimage eccentricity could significantly deviate from zero (i.e., a perfectly foveal location) due to eye movements during afterimage induction. Importantly, afterimage eccentricity had a significant effect on the afterimage’s perceptual null location (PNL) after the saccade, when estimated by both logistic (Figure B2b) and Gaussian (Figure B2c) function.

We modeled afterimage movement to be contingent upon eye movement. Specifically, we defined afterimage position *x_i_* as

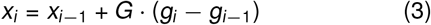

where variable *G* is the afterimage movement gain. After correcting for the varying retinal eccentricity of afterimages (i.e., the offset between inducer stimulus and median gaze position while the inducer was presented) across observers and trials (Figure B2), that was performed by merely adding the estimated eccentricity in each trial to the first recorded gaze position *g*_0_, the starting value *x*_0_ was set equal to *g*_0_. This crucial parameter was fitted to observers’ localization responses by assuming a simple response mechanism (Figure 2b): The model’s localization response *R_i_* depended on *x_i_* relative to the corresponding probe location *p*_*i*−1_ as follows:

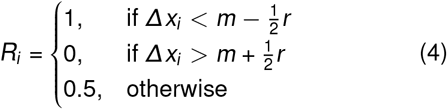

where *Δx_i_* = *x_i_* − *p*_*i*−1_,*m* denotes the response bias, and *r* denotes the range in which the model would report that the afterimage is directly at the probe location. Fitting of the parameters *G, m*, and *r* was performed using a grid-search approach (*G*: [0.8, 1.2] with step size 0.01; *m*: [− 60, 60] pixels with step size 2; *r* : [1, 100] pixels with step size 2) that selected the parameter combination yielding the smallest mean absolute error and the highest classification accuracy across all trials in a given condition for a given observer.

#### Modeling of the inverse relationship

To describe a system that could give rise to afterimage movement, we adapted and expanded an information processing structure proposed by Mittelstaedt (1990). A saccade target with an eccentricity of *z* has to be localized by a (cyclopean and one-dimensional) oculomotor system equipped with motor and optic subunits. The motor subunit receives the motor command *y_c_* and rotates the eye by angle *y* according to

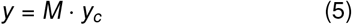

where *M* is the motor gain. The optic subunit receives signal *x* and transforms it to visual space by applying the visual gain *O* as follows:

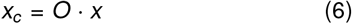

The egocentric location of the target *i* is computed by combining the visual signal and the efference-copy signal given by the forward (motor-to-visual) model:

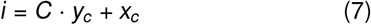

where *C* is the gain of the forward model. Mittelstaedt (1990) shows that, thanks to visual feedback provided by the contingency *x* = *z* − *y*, the equation can be rewritten to

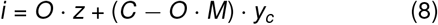

and that, in principle, localization can be achieved “without any extraretinal information about eye position” (Mittelstaedt, 1994, p. 269). However, it remains unclear how the motor command *y_c_* is determined. For that purpose, an inverse model (cf. Masselink & Lappe, 2021) transforms visual coordinates into a motor command capable of foveating *z* by applying the inverse gain *I*:

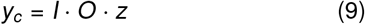

With this assumption *i* can be computed according to

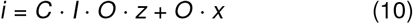

where the first term is the feedforward prediction of the visual consequences of the saccade command. In our case of afterimage localization, the afterimage has a constant (and after inducer-offset correction negligible) retinal eccentricity, reducing its perceived location to

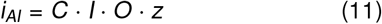

and thereby revealing that *C* and *I* should have an inverse relationship to match afterimage location *i_AI_* with saccade-target location *z*.

## Data and code availability

All preprocessed data, preprocessing and analysis scripts, as well as modeling results can be found at https://osf.io/9egna/. All raw experimental data is furthermore made available at https://osf.io/q3zyh/.

## Acknowledgements

R.S., M.R., and J.R. were funded by the Deutsche Forschungsgemeinschaft (DFG, German Research Foundation) under Germany’s Excellence Strategy – EXC 2002/1 “Science of Intelligence” – project number 390523135. R.S. was supported by the Italian Ministry for Universities and Research (MUR) and the European Union within the Next Generation EU framework (grant ‘T-GAZE’, MSCA_0000027_FIS02) during the analysis and writing stages of the study. M.R. has received funding from the European Research Council (ERC) under the European Union’s Horizon 2020 research and innovation programme (grant agreement No 865715) as well as from the Heisenberg Programme of the DFG (grants RO3579/8-1 and RO3579/12-1).

## Author contributions

R.S. conceived study and built equipment. R.S. implemented and conducted experiment. R.S. analyzed data and performed modeling. R.S. drafted the manuscript, and T.S., J.R. and M.R. provided critical revisions.

## Appendix A Pre-experiment

In the pre-test, observers reported the perceived direction of systematically varied and strictly intra-saccadic inward or outward displacement of the initial saccade target (Figure A1a), estimating not only their discrimination performance but also the target’s perceptual null location (PNL). The task procedure of the pre-test is described in detail in section Pre-experiment. Note that all target manipulations were executed strictly during saccades (Figure A1b) in all conditions (normal jump: *M* = 15.1 ms, *SD* = 0.24; pre-sacc. blanking: *M* = 15.0 ms, *SD* = 0.29; post-sacc. blanking: *M* = 15.0 ms, *SD* = 0.28; pre- and post-sacc. blanking: *M* = 15.0 ms, *SD* = 0.29). Psychometric functions were fitted as described in Analyses, while an iterative procedure that systematically varied starting parameters ensured that fits converged. Individual parameters of logistic fits and group averages (estimated non-parametrically from the average psychometric functions shown in Figure A1c,e using the R package ‘modelfree’; Marin-Franch, Zychaluk, & Foster, 2023) are shown in Figure A1d. Repeated-measures ANOVAs revealed significant differences between presentation conditions were found for all estimated parameters of the psychometric function (*α*; *F* (3, 33) = 14.27, *η*^2^ = 0.46, *p* < .001, *p_GG_ <* .001; *β*; *F* (3, 33) = 12.55, *η*^2^ = 0.45, *p* < .001, *p_GG_* = .001; *λ*; *F* (3, 33) = 4.25, *η*^2^ = 0.07, *p* = .012, *p_GG_* = .035). Finally, a significant negative relationship between observers’ mean primary saccade amplitude and the Gaussian PNL was found (*β* = −0.54, *t* = −4.04, *p* = .002), whose slope did not significantly different in any of the conditions.

Notably, we first replicated the recognized finding that introducing post-saccadic but not pre-saccadic blanking intervals improved trans-saccadic displacement discrimination performance (Figure A1c–e; cf. Born, 2019; Deubel et al., 1996), corroborating the role of blanking in relieving saccadic suppression of displacement (Bridgeman et al., 1975). Results revealed a considerable bias to report inward displacements that manifested itself not only in estimated upper asymptotes as low as 60% but also in PNLs of up to 2 degrees of visual angle (dva). In other words, saccade targets had to be shifted considerably in the direction of the saccade to be perceived as stationary across saccades – a tendency that was strongest in the post-saccadic blanking condition (Figure A1d). This significant saccade direction-contingent shift was likely a consequence of the target’s visual persistence (for a definition and comparison with afterimages, see Di Lollo, Clark, & Hogben, 1988) specific to our experimental setup situated in total darkness: When targets were extinguished briefly before or, as in the post-saccadic blanking condition, even during saccades, the short period of persistence created the impression that targets moved along with the executed eye movement. While computational studies have previously suggested that this effect could be the cause of the phenomenon of peri-saccadic flash mislocalization (Pola, 2004; Teichert et al., 2010), our data provides tentative evidence that visual persistence could indeed cause an illusory shift of visual targets in the direction of the impending saccade.

**Figure A1.**
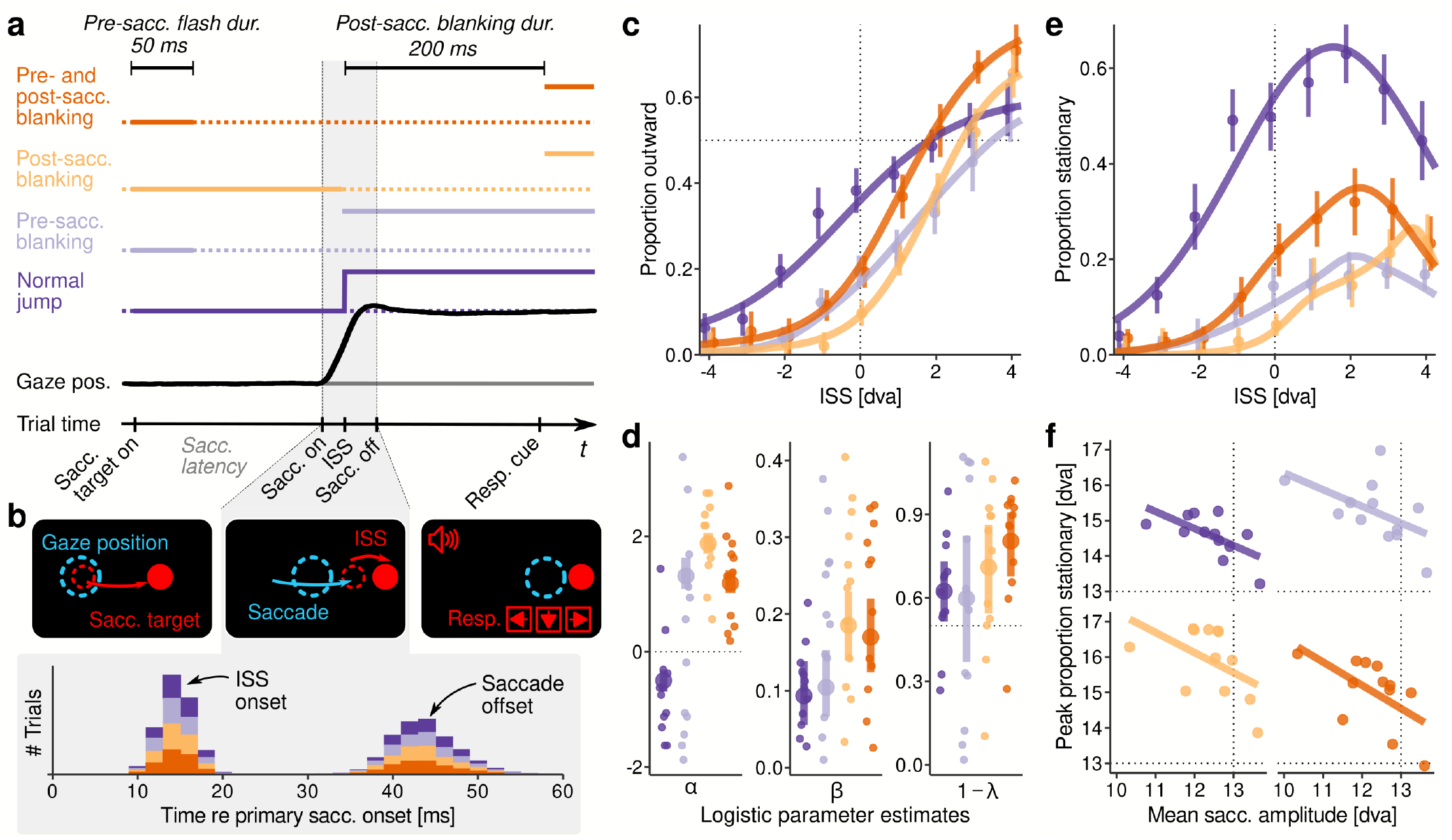
Trans-saccadic displacement judgments in total darkness. **a** Illustration of the temporal structure of the four presentation conditions used to display the intra-saccadic displacement (ISS). **b** Illustration of the spatial structure of the stimulus display (without any blanking) before, during, and after saccade execution. Lower panel shows the distribution of ISS timing relative to physical saccade onset (left histogram) with reference to the distribution of saccade duration (right histogram). **c** Average logistic functions approximating the proportion of ‘outward’ judgments as a function of ISS size. The color scheme follow the definitions in panel a. **d** Logistic-function parameter estimates (γ not shown) for individual observers and group averages (±SEM), plotted by presentation conditions. **e** Average Gaussian function approximating the proportion of ‘stationary’ responses as a function of ISS size. **f** Mean parameters of fitted Gaussian functions as function of average saccade amplitude in displacement-judgment trials, plotted separately for each presentation condition outlined in panel a.

## Appendix B Main experiment

### Evaluation of apparatus

To evaluate how well TTL triggers corresponded with LED activity, we simultaneously measured trigger and photodiode currents. We found that they were most strongly correlated at zero lag (Figure B1b), suggesting that triggers indeed reported physical stimulus onsets with high fidelity. Finally, using a ColorCAL MKII Colorimeter (Cambridge Research Systems, Rochester, Kent, UK), we ascertained the luminance range and the linear gamma of the utilized LEDs (Figure B1c), as well as the specifications of the presented stimulus colors (see Stimuli).

**Figure B1.**
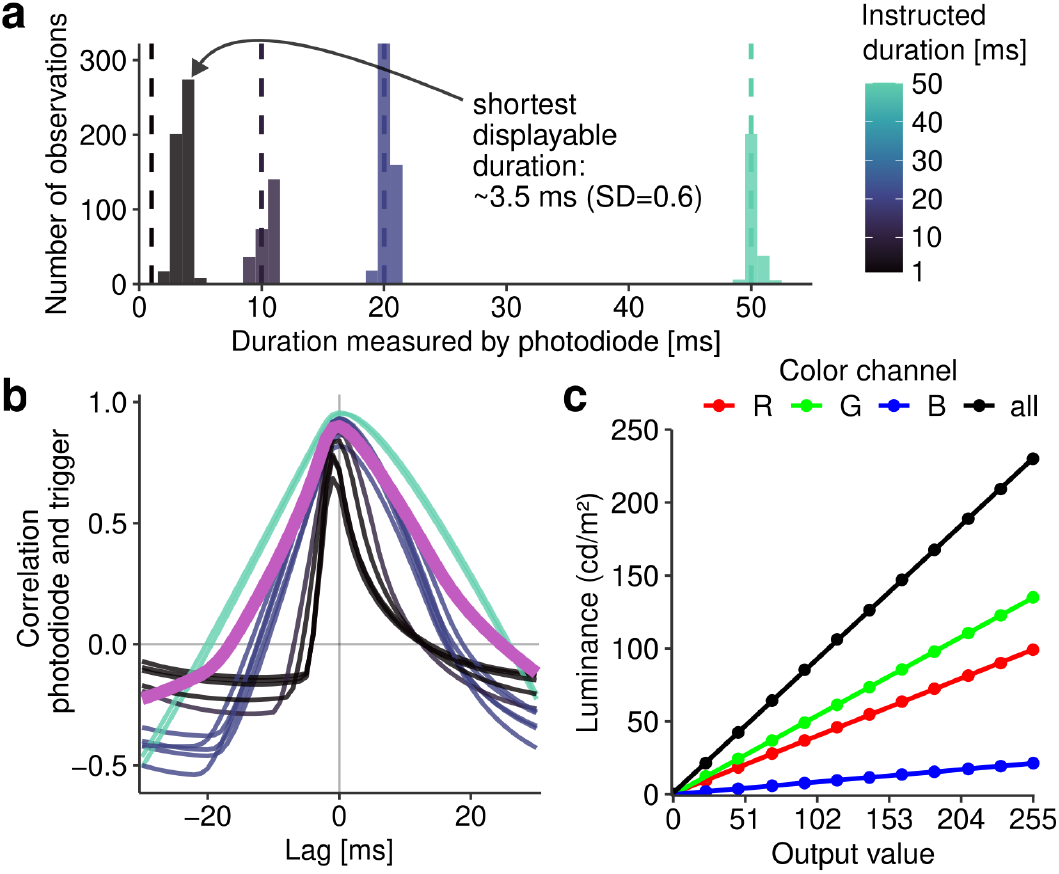
Evaluation of the custom-built LED screen’s properties. **a** Photodiode measurements (sampled at 1000 Hz) of LED on-durations made for four different instructed LED activation durations (vertical dashed lines). **b** Cross-correlation analysis performed on photodiode and trigger signals. The thick purple line indicates the mean across all tested activation durations. **c** Luminance of the LEDs’ RGB color channels, measured across the entire range of output values. Straight lines indicate linear fits of the measurements.

### Estimation of afterimage eccentricity

The retinal eccentricity of afterimages was computed by the offset between the position of the inducer pulse and the median gaze position during inducer display. Figure B2a shows partly considerable and highly individual horizontal eccentricity of afterimages for individual observers. Evidence that afterimage eccentricity had a major impact on perceived afterimage position after the eye movement is provided by Figure B2b-c: Mean afterimage eccentricity significantly predicted the logistic afterimage PNL after the execution of eye movements (*β* = 0.40, *t* = 2.96, *p* = .008) and similar slopes, yet not significant, were estimated for the Gaussian afterimage PNL (*β* = 0.34, *t* = 1.94, *p* = .066). Both of the slopes were similar across saccade directions (logistic: *β* = −0.06, *t* = −0.47, *p* = .644; Gaussian: *β* = −0.001, *t* = −0.004, *p* = .997). If correction of afterimage eccentricity were not performed, then group-level averages of afterimage PNLs may be misinterpreted (see transparent lines in Figure 2d that show psychometric functions without correction). In fact, the correction significantly improved model fits increasing classification accuracy by on average 1.6% (*F* (1, 11) = 18.2, *η*^2^ = 0.05, *p* = .001).

**Figure B2.**
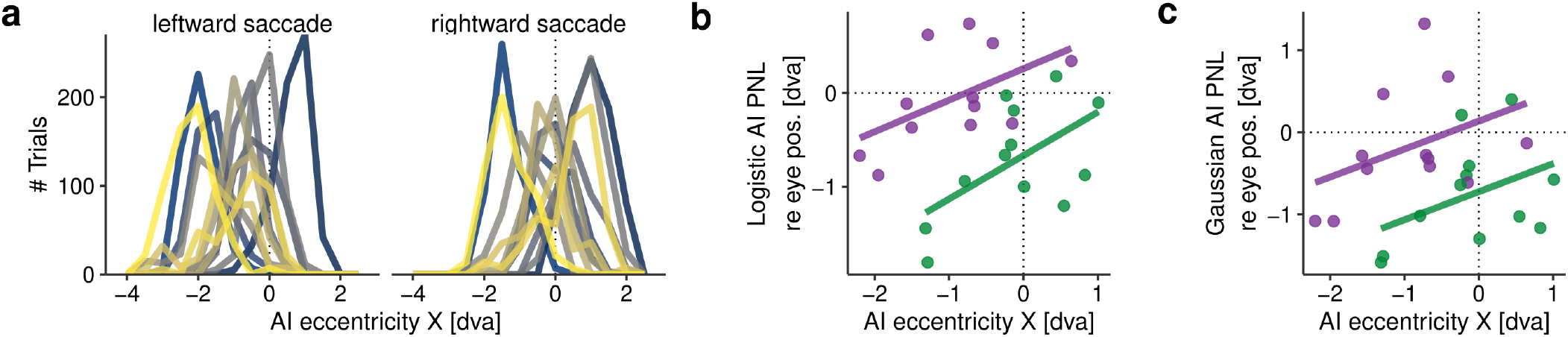
Retinal eccentricity of afterimages. **a** Distribution of horizontal afterimage eccentricity, shown for individual observers and leftward and rightward saccades, respectively. **b-c** Relationship between mean afterimage eccentricity and (post-saccadic) afterimage PNL, as estimated by the logistic (panel b) and Gaussian (panel c) models, using the same color conventions as in Figure 2 (green: rightward, purple: leftward).

### Primary and secondary saccades

First, we plotted observers’ *G* parameters, estimated in Session 1 (Figure 6c), as a function of their primary saccade gain, revealing that one observer (Observer 4) who made drastically hypometric saccades, while her or his estimated gain *G* was normal (Figure B3a). To understand this curious deviation, we extracted observers’ secondary saccade metrics and found that Observer 4 made secondary saccades of a sufficient gain to fully compensate for hypometric primary saccades (Figure B3b). These secondary saccades were extremely frequent and were initiated with very short latencies (Figure B3c), suggesting they may have been programmed as part of the initial saccadic motor command to reach the target. The reason why gains were normal for Observer 4 is owing to the fact that the Gain Model accounted for all eye movements from the start of the primary saccade to the presentation of the first localization probe.

**Figure B3.**
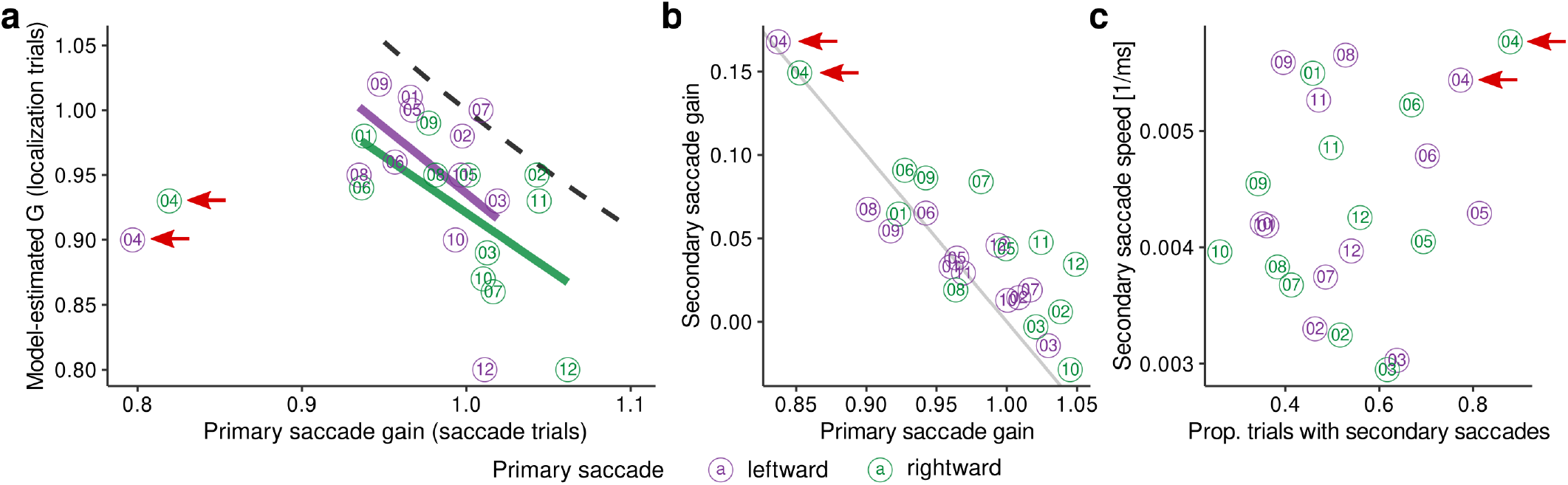
Analysis of observers’ saccade strategies. **a** Model-estimated gain parameters as a function of primary saccade gain for individual observers (indicated by number) for rightward (green) and leftward (purple) saccades, respectively. Black dashed line indicates the inverse relationship. Data points of Observer 4 are highlighted by arrows. **b** Secondary saccade gain as a function of primary saccade gain. Dotted line indicates a linear relationship with a slope of −1 and intercept of 1. **c** Relationship between proportion of trials in which secondary saccades were made and the speed with which they were initiated after primary saccade offset (i.e., the inverse of secondary saccade latency).

## Notes

We have no conflict of interest to disclose.

https://osf.io/9egna/

### Competing Interest Statement

The authors have declared no competing interest.

### Summary of Updates

Shortening of abstract, additional analysis to validate the assumption of linear gains, clarifications and edits for readability in the main text

